# Engineered endosymbionts capable of directing mammalian cell gene expression

**DOI:** 10.1101/2021.10.05.463266

**Authors:** Cody S. Madsen, Ashley V. Makela, Emily M. Greeson, Jonathan W. Hardy, Christopher H. Contag

**Affiliations:** Department of Biomedical Engineering, Michigan State University, East Lansing, MI, USA; Department of Microbiology and Molecular Genetics, Michigan State University, East Lansing, MI, USA; Institute for Quantitative Health Science and Engineering, Michigan State University, East Lansing, MI, USA

**Keywords:** Engineered endosymbiont, intracellular bacterial therapeutic, immune modulation, bacteriotherapy, nuclear localization

## Abstract

Modular methods for directing mammalian gene expression would enable advances in tissue regeneration, enhance cell-based therapeutics and improve modulation of immune responses. To address this challenge, engineered endosymbionts (EES) that escape endosomal destruction, reside in the cytoplasm of mammalian cells, and secrete proteins that are transported to the nucleus to control host cell gene expression were developed. Microscopy confirmed that EES escape phagosomes, replicate within the cytoplasm, and can secrete reporter proteins into the cytoplasm that were then transported to the nucleus. Synthetic operons encoding the mammalian transcription factors, *Stat-1* and *Klf6* or *Klf4* and *Gata-3* were recombined into the EES genome. Using controlled induction, these EES were shown to direct gene expression in J774A.1 macrophage/monocyte cells and modulate the host cell fates. Expressing mammalian transcription factors from engineered intracellular bacteria as endosymbionts comprises a new tool for directing host cell gene expression for therapeutic and research purposes.

**Graphical abstract:** 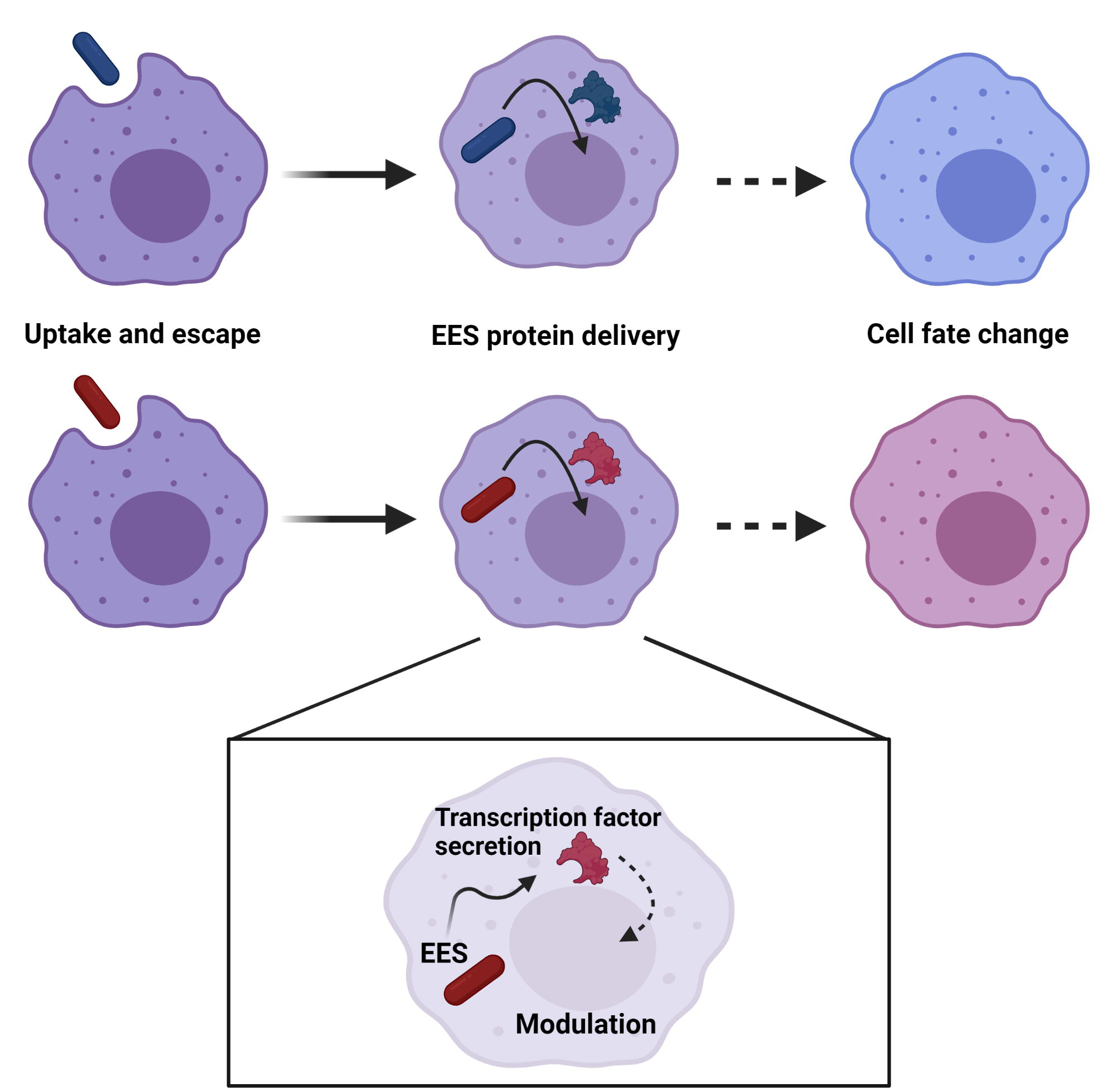

## Introduction

The currently accepted theory of the origin of eukaryotic cells postulates that at an early stage of evolution, there was a close and long-term biological interaction (symbiosis) between separate single-cell organisms (Sagan, 1967). Free-living bacteria (prokaryotes) were engulfed by other primitive cells as endosymbionts, and through elimination of unneeded duplicative functions and selection of mutually beneficial processes, this eventually created stable internal structures (organelles) that provide specific functions not encoded in the genome of the host cell (Sagan, 1967). The genesis of organelles such as mitochondria and chloroplasts led to the evolution of eukaryotic cells and multicellular organisms and is referred to as endosymbiogenesis (Corsaro et al., 1999). This concept inspired this study’s construction of engineered endosymbionts (**EES**) to direct mammalian cell fates and function. The term EES refers to bacteria that remain viable in the cytoplasm of mammalian cells over extended periods of time, either naturally or through modification, and are engineered to produce mammalian modulators (proteins, metabolites or nucleic acids) that can redirect host cell biology.

The concept of using endosymbionts for modifying eukaryotic organisms has been utilized for a decade in insect hosts. *Wolbachia spp*. is a model endosymbiont that lives symbiotically within mosquitoes and naturally blocks the transmission of dengue and Zika virus by the mosquito species *Aedes aegypti* (Jiggins, 2017). Release of mosquitoes infected with *Wolbachia spp*. into the environment is a strategy to prevent transmission of viruses (Flores and O’Neill, 2018; Jiggins, 2017). Research on natural endosymbionts of invertebrates has revealed benefits of the symbiotic relationships (Chao et al., 2021; Ip et al., 2021; Izraeli et al., 2021; Kennedy et al., 2007; Shi et al., 2021; Xu et al., 2021; Yang et al., 2021).

A recent study has investigated mimicking mitochondria by engineering a bacterium to act as an EES (Mehta et al., 2018). In this study, *Escherichia coli* (*E. coli*) was engineered to exist in the cytoplasm of *Saccharomyces cerevisiae* (*S. cerevisiae*) and supply adenosine triphosphate (ATP) as a replacement for deficient mitochondria of the yeast host cell. In return, *S. cerevisiae* produced an essential vitamin for *E. coli* to create a synthetic, or engineered, symbiotic relationship. This relationship was stable over several generations of *S. cerevisiae*, revealing that an extracellular bacterium could be engineered to live inside a eukaryotic cell in a dependent relationship. This EES was developed to advance our understanding of endosymbiosis. In another study, extracellular bacteria were investigated as a means of delivering proteins to mammalian cells for the purpose of altering their behavior (Jin et al., 2018). In this case, *Pseudomonas aeruginosa* delivered transcription factors to induced pluripotent stem cells (iPSCs) by a type III secretion system, which differentiated iPSCs into cardiomyocytes (Jin et al., 2018). These studies reveal an opportunity for the development of a stable, non-pathogenic EES capable of delivering transcription factors to mammalian cells and altering cellular fates and functions.

A target for the development and use of stable, non-pathogenic EES are phagocytic immune cells. The immune system is delicately balanced in mammalian systems, both recognizing self and building tissues, and defending against disease and foreign invaders capable of damage. However, loss of immunological homeostasis (*i*.*e*. balance) may contribute to disease progression, and this process has been extensively studied in the context of macrophage function (Ginhoux and Guilliams, 2016; Kim et al., 2019). Macrophages are an abundant cell throughout the body, playing important roles in host immunity, tumor progression and modulation of the host response (Chu et al., 2020; Hume, 2008; Wynn et al., 2013). They are commonly recruited by stimuli in response to inflammation and contribute to progression of associated pathologies, thus representing a significant component of the inflammatory microenvironment and the resulting outcome (Ginhoux and Guilliams, 2016; Kim et al., 2019), and as a target for engineering. As a first line of defense, macrophages are phagocytic, acting to take up and destroy foreign invaders or damaged cells (Epelman et al., 2014), which would include extracellular EES. Macrophages are also remarkably plastic, with the ability to switch phenotypes and alternate between synthesizing proinflammatory or anti-inflammatory signals (Lavin et al., 2014). Their function is influenced by the microenvironment in which they reside, where they respond to a variety of signals and stimuli. These mechanisms are often co-opted by tumors to create anti-inflammatory microenvironments (Epelman et al., 2014; Pawelek et al., 2006). There are broadly two categories of macrophages, associated with distinct and opposing functions once stimulated from resting state (M0). Pro-inflammatory macrophages (M1) act in a fashion to destroy pathogens, including tumor cells, and anti-inflammatory (M2) macrophages decrease inflammation, support angiogenesis and promote tissue remodeling and repair (Moreira and Hogaboam, 2011; Mosser and Edwards, 2008; Shaughnessy and Swanson, 2007; Zhang and Mosser, 2008). Inflammation states can be distinguished by changes in cell surface markers (*e*.*g*. cluster of differentiation (CD)86; M1 or CD206; M2) or through profiling of cytokines (Mantovani et al., 2004).

Commonly, the phenotypic change, or polarization of macrophages that is associated with disease can act to either induce or suppress the progression of pathology (Chen et al., 2020). Therefore, macrophages may prove to be a key cell for molecular therapies directed at modifying cellular functions, since these cells are present within injured, damaged and malignant tissues and can be modulated to switch phenotypes to alter the disease course (Martinez et al., 2008; Porcheray et al., 2005). The use of a living organism, such as an engineered bacterium residing inside a mammalian cell, could provide a means of control. Precedence exists for using bacteria to impact mammalian cell physiology for therapeutic approaches, known as bacteriotherapy, and advances in this field support the development of EES for cellular control. Bacille Calmette-Guerin (BCG, the *Mycobacterium bovis* strain used as a tuberculosis vaccine) bacteriotherapy has become standard of care for bladder cancer treatment, and many other clinical applications for bacteriotherapy are being devised and tested, including for the treatment of other cancers (Laliani et al., 2020; Sawant et al., 2020; Sedighi et al., 2019; Soleimanpour et al., 2020; Yaghoubi et al., 2019, 2020a; Zhang et al., 2020). Bacterial systems, including EES, represent a potentially powerful means of exploiting prokaryotic versatility for therapeutic purposes.

The concept of bacterial endosymbiont-driven modulation of mammalian cells begins by developing a bacterial endosymbiont to control mammalian cell fates and function. *Bacillus subtilis* expressing listeriolysin O (LLO) is an engineered intracellular bacterium (Bielecki et al., 1990) that was created from *B. subtilis* ZB307 (a derivative of *B. subtilis* strain 168) (Zuber and Losick, 1987). LLO lyses the phagocytic vacuole, releasing ingested bacteria into the cytosol. In this strain, the *hylA* gene that encodes LLO from *Listeria monocytogenes*, was placed under control of an isopropyl β-D-1-thiogalactopyranoside (IPTG) inducible promoter and inserted into the genome of *B. subtilis*, allowing the bacteria to escape phagosomes in mammalian cells when *hylA* is induced (Bielecki et al., 1990; Portnoy et al., 1992). Since *B. subtilis* LLO can access the cytoplasm and does not have a lipopolysaccharide (LPS) mediated immune response (Travassos et al., 2004), *B. subtilis* LLO was chosen as the chassis organism to create an EES. Additionally, *B. subtilis* is a non-pathogenic, Gram-positive, soil bacterium that respires as a facultative anaerobe thus making it possible to respire in the cytoplasm of the host cell (Gallegos-Monterrosa et al., 2016; Nakano and Zuber, 1998). This bacterium has been classically used for secreting complex proteins into the surrounding extracellular space through the general secretory (Sec) and twin-arginine translocation (Tat) pathways (Kolkman et al., 2008). *B. subtilis* has been well characterized to the point of full genome annotation, databases (*e*.*g*. BsubCyc database) have been developed for metabolism analysis and protein production (Barbe et al., 2009), and contain multiple inducible systems including several sugar-regulated inducible systems (Vavrová et al., 2010). *B. subtilis* contains a promoter-regulator inducible system that is sensitive to D-mannose. D-mannose has been shown to be actively transported inside of mammalian cells which provides a second inducible system to regulate EES gene expression (Panneerselvam and Freeze, 1996; Sun and Altenbuchner, 2010). A goal of this study was to demonstrate the functionality of EES, within the cytoplasm of live mammalian cells. Initially, the EES were designed to secrete β-galactosidase (β-gal) from the entire *lacZ* gene via the Tat secretion pathway through signaling of the PhoD secretion peptide (Kolkman et al., 2008; Tjalsma et al., 2000). Nuclear trafficking was accomplished using the simian virus (SV) 40 nuclear localization signal (NLS) that directed *E. coli* β-gal expressed from mammalian ribosomes to the nucleus previously (Kalderon et al., 1984).

Here the EES is used to modulate mammalian cell gene expression by engineering operons to express mammalian transcription factors that are secreted from the EES and delivered to the nucleus of mammalian cells (macrophages) to control gene expression (Figure 1, Figure S1). One operon encodes the transcription factors signal transducer and activator of transcription 1 (STAT-1) and Krüppel-like factor 6 (KLF6), and the second encodes Krüppel-like factor 4 (KLF4) and GATA binding protein 3 (GATA-3) (Biswas et al., 2006; Date et al., 2014; Li et al., 2018; Liao et al., 2011; Yang et al., 2018). These transcription factors have been shown to promote transition from resting state (M0) to alternative, M1- or M2-shifted phenotypes, respectively. The activity of STAT-1 is directly controlled by the presence of pro-inflammatory IFN-γ and cytokines while KLF6 has been shown to be upregulated in M1 polarized macrophages. In parallel, the activity of KLF4 is directly stimulated by the signal cascade from the cytokine interleukin 4 (IL-4) which promotes M2 polarization and GATA-3 is highly upregulated in M2 macrophages (Biswas et al., 2006; Date et al., 2014; Li et al., 2018; Liao et al., 2011; Yang et al., 2018). Therefore, each operon could provide a directed approach towards the respective polarization while also demonstrating EES modulation of gene expression.

**Figure 1.**
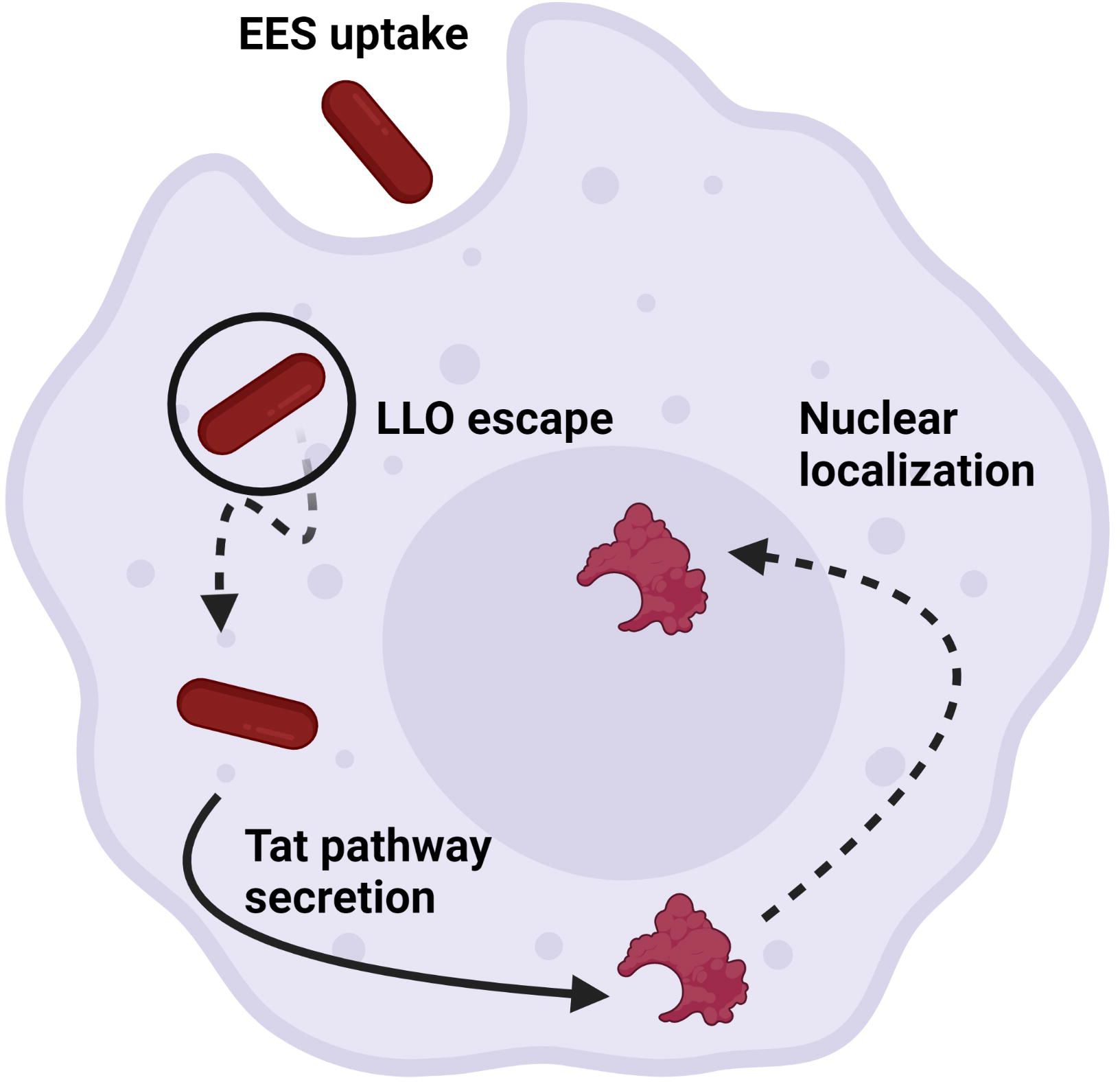
EES as a means of controlling gene expression in mammalian host cells. The EES enter phagocytic mammalian host cells and escape the phagosome using the LLO protein. The EES then secrete a reporter protein or transcription factor into the cytoplasm through the Tat pathway followed by localization to the host cell nuclei.

## Results

### EES escape from phagosomes into cytoplasm of J774A.1 cells

Confocal microscopy confirmed the escape of the EES from phagosomes after uptake into J774A.1 cells. In Figure 2, EES (magenta) localization was compared to LAMP-1 (Kortebi et al., 2017) positive structures (phagosomes, red) in J774A.1 cells (green) with and without IPTG induction of LLO expression (+IPTG and -IPTG). Without IPTG induction, very few EES were observed in the J774A.1 cultures, and many regions of punctate signal, commonly within LAMP-1 positive regions were seen (Figure 2, zoom-solid line; Movie S2). Localization of the EES stain (magenta) within the phagosome structures (red) indicated that the EES were degraded in the absence of induced LLO expression. Escape into the cytoplasm only occurred when the *hylA* gene (LLO) was induced (Movie S2). The IPTG-inducible promoter is known to allow some transcription in the absence of IPTG; this likely resulted in the few EES that were observed in the cytoplasm that were not entrapped in phagosomes (Figure 2; -IPTG, zoom-dotted line). In contrast, many EES were present throughout the mammalian cells and did not colocalize with LAMP-1 staining, *i*.*e*. were in the cytoplasm, when LLO was induced with IPTG. Within the macrophages, there were both EES which had not yet escaped the phagosome (Figure 2; +IPTG, white arrowhead) and EES which had escaped and were present in the cytoplasm (Figure 2; +IPTG, zoom-solid line). 3D analysis using z-stack data confirmed presence of EES throughout the depth of J774A.1 cells that were in the cytoplasm and not associated with LAMP-1 positive structures (Movie S1). Confocal imaging confirmed the presence of EES in the cytoplasm as previously reported using other methods (Bielecki et al., 1990).

**Figure 2.**
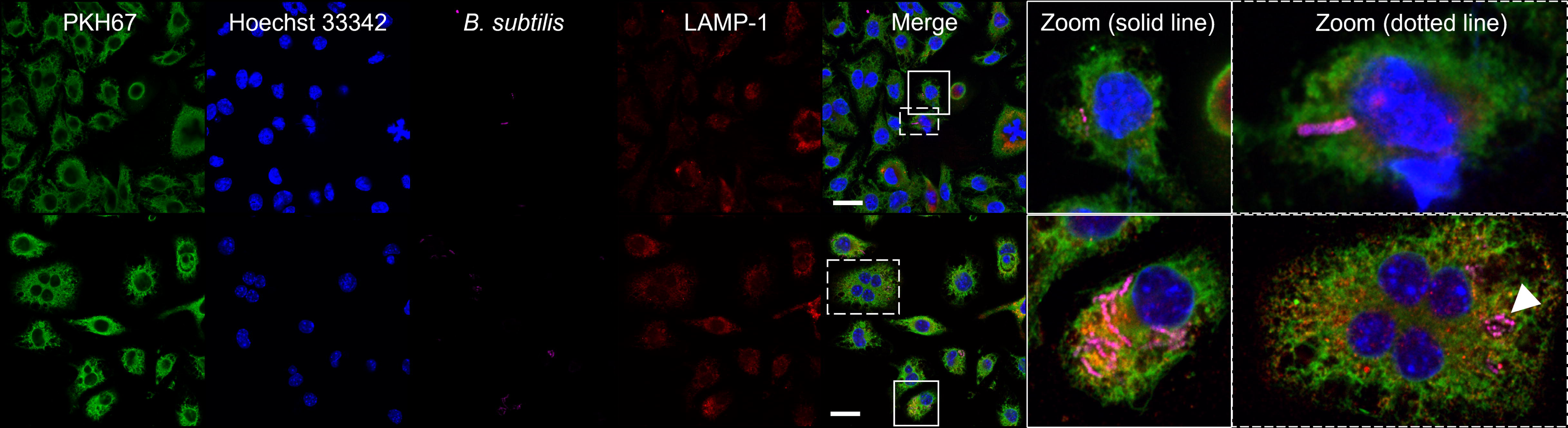
Confirmation of EES phagosomal escape into the cytoplasm of J774A.1 cells by confocal imaging. Confocal imaging of EES in J774A.1 cells visualized the PKH67 membrane stain (green), nuclear Hoechst 33342 (blue), anti-*B. subtilis* (magenta) and anti-LAMP-1 (red). EES were introduced to J774A.1 cells and treated without (top) and with IPTG (bottom). Merged and zoomed panels display EES escape when LLO was induced with IPTG; +IPTG green channel intensity was decreased, and magenta channel intensity increased in zoomed images to visualize bacteria. Scale bars = 20 µm.

### Viability of EES and J774A.1 cells during EES cytoplasmic persistence

J774A.1 host cell viability was assessed using an MTS cell proliferation assay. At 1 hour (hr) post EES addition, there was no significant change in host cell viability in all treatment conditions except at the highest multiplicity of infection (MOI, 50:1; Figure 3A). At 4 hr post EES addition, both the 25:1 MOI and 50:1 MOI conditions had significant loss in host cell viability (Figure 3A). The same trend was observed after staining host cells with a viability dye, after EES addition, and assessing cell viability with flow cytometry (Figure S2). Live cell imaging demonstrated that the intracellular EES replicated in the host cell after phagosomal escape (Figure 3B). Increasing numbers of EES were visualized over time, post co-incubation with a 10:1 MOI. Brightfield imaging visualized the J774A.1 cells, with the EES (magenta) within the cytoplasm (Figure 3B). The cells were imaged at 1 hr and 2.5 hr post EES addition. Zoomed in regions indicate each EES doubled twice during the two time points from 3 EES at 1 hr to 12 at 2.5 hr. Additionally, the reproducibility of the interaction between EES and host cell was quantified. The number of cells containing EES and the EES per cell were determined under the same conditions as in the MTS assay (Table S1). Quantification of EES after treatment with an MOI of 10:1 confirmed the replication of the EES across the population (Table S1). At each MOI tested, approximately 50% of the EES entered the host cells, and using MOIs of 25:1 or 50:1, nearly 100% of the host cells contained EES (Table S1). Therefore, combining the results from MTS and live cell quantification, 25:1 MOI was chosen for the EES delivering protein to the nuclei of the host cells.

**Figure 3.**
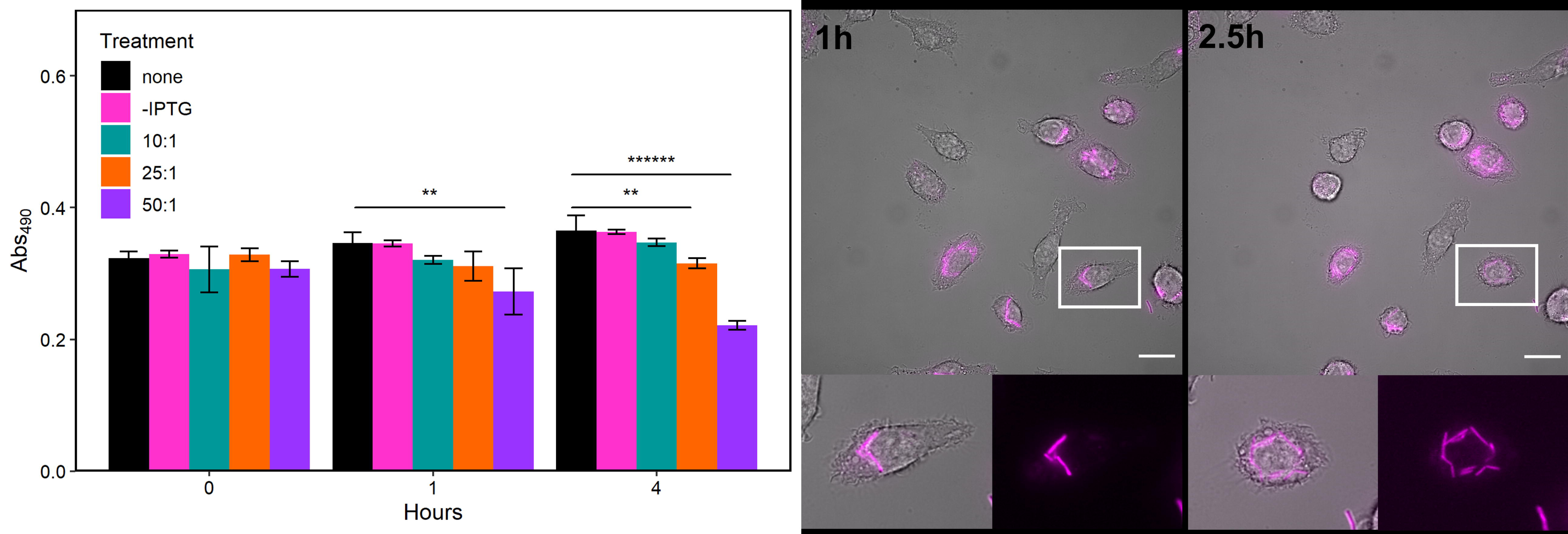
Host cell viability and replication of EES in the cytoplasm of J774A.1 cells. An MTS assay quantified viability of the J774A.1 cells at multiple time points after treatment with EES under various conditions (left). Live cell microscopy identified EES replication in the host cell at 1 hr and 2.5 hr post EES addition (right). J774A.1 cells were visualized in brightfield, and EES (magenta); zoomed images reveal EES replication in the cytoplasm. Plotted data is mean ± SD; **p<0.01, ******p<0.000001. Scale bars = 20 µm.

### Inducible EES secretes β-gal with delivery to the nuclei of J774A.1 cells

The production of β-gal by EES-*lacZ*, with and without a nuclear localization signal (NLS; EES-*lacZ*-NLS and EES-*lacZ*-no NLS), was studied with IPTG or mannose control. Localization of the reporter protein to the J774A.1 nuclei was confirmed after coincubation with EES and end-point fluorescence microscopy (Figure 4, S3). The presence of β-gal in the nuclei of host cells was determined by measuring the fluorescent signal to noise ratio (SNR) of the nuclei compared to background signals. Nuclear SNR of cells with EES-*lacZ*-NLS, after induction with mannose for 3 hr, was 3.5-fold higher than that of J774A.1 cells only or cells incubated with β-gal protein obtained from sterile filtered supernatant (1 mL) from EES*-lacZ-*NLS cultures after 16 hr of induction with mannose (Figure 4). Additionally, nuclear SNR of cells incubated with EES-*lacZ*-NLS and mannose was significantly higher than without mannose induction and that of EES-*lacZ*-no NLS (Figure 4). The mannose-inducible system using 3 hr incubation showed significantly increased amounts of β-gal in the nucleus over the IPTG-inducible system and provided a dual inducible system for control in the EES (Figure S3). Further, a high concentration of gentamicin was added after 3 hr of induced β-gal production to rescue the J774A.1 cells from overgrowth of the EES and allow for another 21 hr of trafficking of protein to the nucleus. Nuclear β-gal SNR of cells exposed to mannose-induced EES-*lacZ*-NLS for 3 hr and then incubated for 21 additional hours (Figure 4) was 5.1-fold higher than that of J774A.1 cells only, or cells incubated with the supernatant of induced EES-*lacZ*-NLS. Nuclear SNR of cells exposed to β-gal trafficking for 24 hr, delivered by the mannose-induced EES-*lacZ*-NLS, was 2-fold higher than with no mannose and 1.5-fold higher than those exposed to the EES-*lacZ*-no NLS strain (Figure 4). Consequently, the mannose-inducible system in the EES was shown to be controlled inside mammalian cells and to provide regulation in delivering protein to the J774A.1 nuclei.

**Figure 4.**
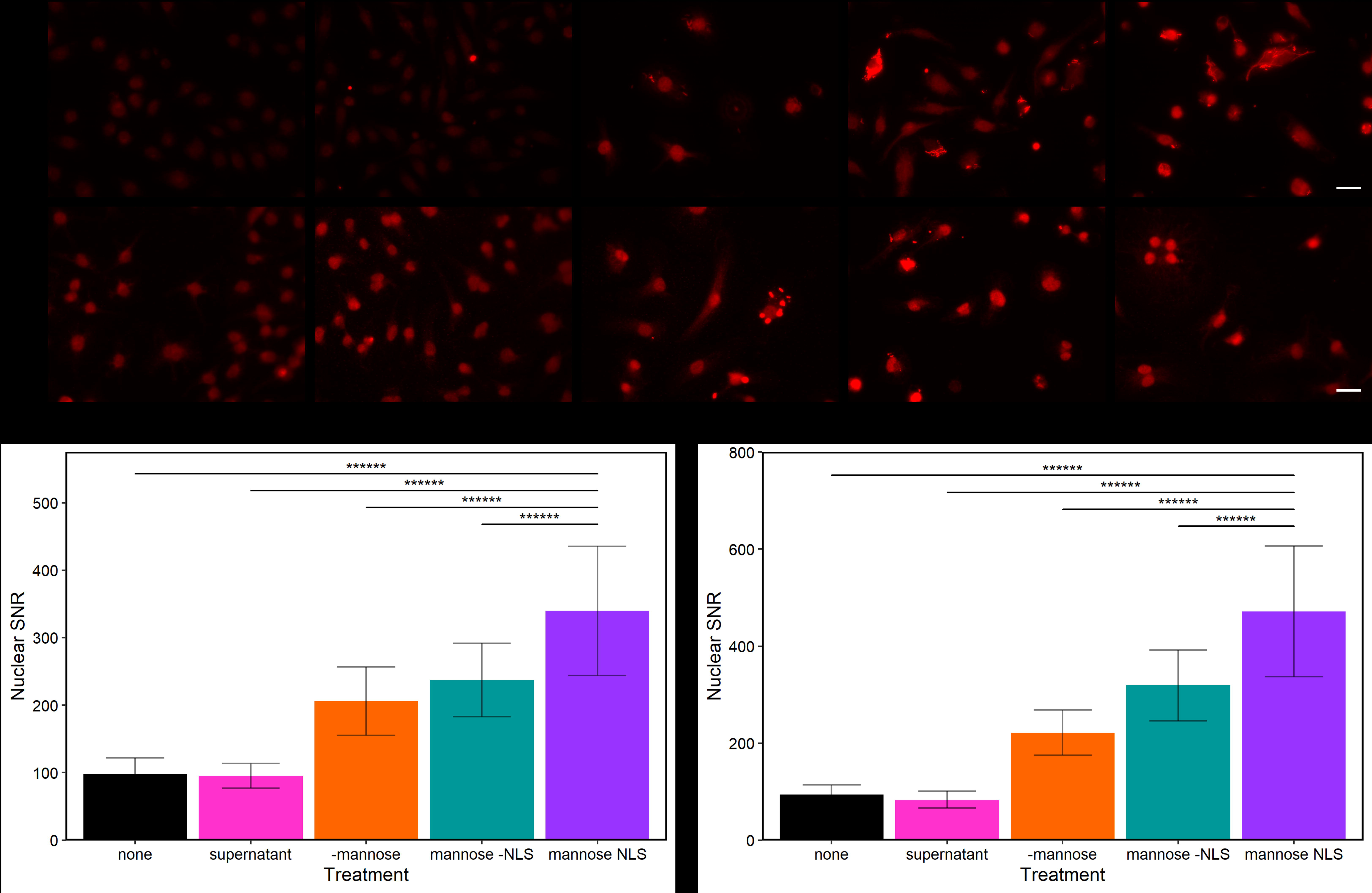
Intracellular localization of EES-*lacZ* secreted β-gal. Fluorescence microscopy (top) and signal to noise ratio (bottom) of nuclei in J774A.1 cells with no EES (none), J774A.1 cells incubated with β-gal collected as supernatant from induced EES-*lacZ*-NLS (supernatant), J774A.1 cells incubated with uninduced EES-*lacZ*-NLS (-mannose), J774A.1 cells incubated with induced EES-*lacZ*-no NLS (mannose -NLS) and J774A.1 cells incubated with induced EES-*lacZ*-NLS (mannose NLS). Plotted data is mean ± SD; ******p<0.000001. Scale bars = 20 µm.

### EES modulation of J77A.1 cell gene expression determined by flow cytometry

Flow cytometry detected differences in the levels of macrophage polarization-associated cell surface markers, CD86 (M1 polarization) and CD206 (M2 polarization), in J774A.1 cells under various conditions, at 24 or 48 hr time points, post incubation (Figure 5, S5). At 24 hr post incubation, there was a significant increase in CD86 expression in J774A.1 cells containing EES-*Stat-1Klf6* (EES-*SK*), which had been induced with mannose (Figure 5). This induction was observed by an increase in CD86 median fluorescence intensity (MFI) when compared to J774A.1 cells with no treatment, indicating gene-specific M1 polarization induced by *Stat-1* and *Klf6*. Additionally, CD86 MFI from this condition was not significantly different than that of the M1-polarized positive control (*p* = 0.5155). CD206 MFI at 24 hr did not identify any differences in any EES-*Klf4Gata-3* (EES-*KG)* condition versus resting J774A.1 cells. At 48 hr, CD86 MFI was increased in both EES and EES-*SK* -mannose versus resting. There was no significant difference in CD206 expression at 48 hr. Immunofluorescence confirmed production of STAT-1 and KLF6 or KLF4 and GATA-3 in cells containing EES-SK or EES-KG, respectively (Figure S4).

**Figure 5.**
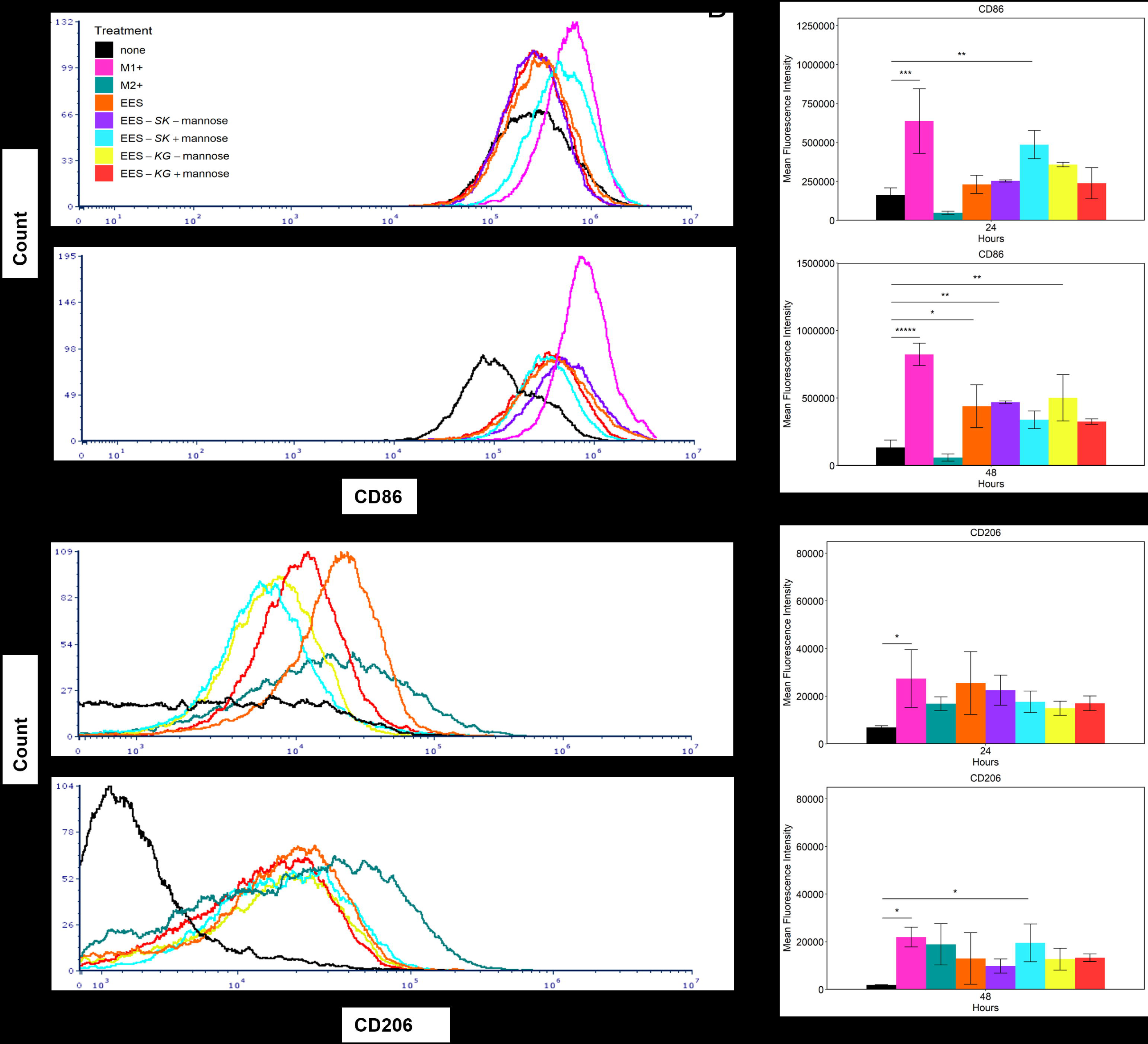
Flow cytometry demonstrating EES can impact J774A.1 cell marker expression. Histograms of median fluorescent intensity (MFI) revealed shifts of CD86 and CD206 expression by EES (A). These data are representative examples, taken as the median value of n = 3. Average CD86 or CD206 MFI is compared between all treatments and statistically significant values compared to ‘none’ treatment are plotted (B). J774A.1 cells were treated with nothing (none), LPS and IFN-γ (M1+), IL-4 and IL-13 (M2+), EES, EES-*SK* with and without mannose (EES-*SK* -mannose, EES-*SK* +mannose) and EES-*KG* with and without mannose (EES-*KG* - mannose, EES-*KG* +mannose) at 24 and 48 hr post initial treatment. Plotted data is mean ± SD; *p<0.05, **p<0.01, ***p<0.001, *****p<0.00001.

J774A.1 cells that contained an EES versus those that did not were analyzed for CD86/CD206 expression (Figure S5). There was no significant difference in the CD86 (*p* = 0.9891) and CD206 (*p* = 0.9987) expression between cells that did and did not contain EES at 24 hr. However, at 48 hr there were differences in these populations. The percent of CD86+ cells was different between cells that did and did not contain EES (*p* = 0.0259), as well as between the different EES (+/- mannose or EES-LLO; p = 0.037). CD206 (*p* = 0.0513) expression was not different in J774A.1 cells that did and did not contain EES. When comparing the entire population, results trended towards shifting the cell population to CD86+ with 24.54 +/- 15.73% of EES-*SK* +mannose, 4.35 +/- 3.66 EES-*SK* -mannose and 20.26 +/- 26.84% EES (Figure S5). This finding was consistent with the CD86 MFI results. There were no differences in the CD206+ population at 48 hr in any EES conditions vs resting J774A.1 cells.

### EES modulation of J77A.1 cell gene expression determined by cytokine profiling

EES-*SK* and EES-*KG* modulated mammalian cell gene expression, compared to that of the native EES strain, as determined by Luminex cytokine profiling (Figure 6, S6). The response patterns were complex, but the distinct effect of the respective genes was clearly observed. All EES strains caused a rapid and significant production in cytokines over the resting state and the positive controls. The most prominent cytokine changes are shown below; 14 of the 17 cytokines in the panel revealed gene-specific distinctions between the EES-*SK* and EES-*KG* strains. Cytokine production was generally higher at 24 hr compared to 48 hr post EES exposure for most affected cytokines, while the significant differences were observed at both time points. Activating the transcription factor operons with mannose dramatically affected cytokine production in J774A.1 cells containing EES-*SK* or EES-*KG*, and distinctions between the EES with the different operons were apparent (Figure 6). Addition of EES-*SK* and EES-*KG* strains to J774A.1 cells led to increased and decreased production of the same cytokines in comparison to EES, respectively, as shown by IL-10 and IL-12p40 (Figure 6A-C). Additionally, both strains were able to downregulate tumor necrosis factor alpha (TNF-α) nearly 20-fold for each operon strain (except EES-*KG* -mannose). EES-*SK* was able to downregulate granulocyte colony stimulating factor (G-CSF) (Figure 6D-E). However, D-mannose alone may have had a significant impact on certain cytokines such as IL-1α and IL-1β; significant results were observed in the treatment conditions when D-mannose was added and caused the same impact on the cytokine (Figure 6F, Figure S6C-D). Production of certain cytokines was not impacted by any treatment condition, including vascular endothelial growth factor (VEGF). Several cytokines were only significantly changed at one of the two time points in certain conditions, including IL-6 and macrophage inflammatory protein-2 (MIP-2/CXCL2) (Figure S6F-G, O-P). Lastly, some cytokines such as eotaxin-1 (CCL11), IFN-γ, and leukemia inhibitory factor (LIF) were not significantly produced by the J774A.1 cells (not shown). MIP-1α production at 24 hr was outside the concentration of the standards even after dilution (also not shown). IL-4 only appeared in the M2+ condition in which it was added. IL-13 exhibited the same trend with minimal expression at 24 hr in the other treatment conditions besides the M2+ (Figure S6H-I). The EES-*SK* and EES-*KG* strains were able to significantly impact mammalian cell gene expression compared to positive controls and the base EES strain, which had a significant impact on cytokine production in J774A.1 cells (Figure 6, S6). Ultimately, flow cytometry measurements demonstrated the EES-*SK* was able to change marker expression similar to that of the M1-positive control and both the EES-*SK* and EES-*KG* impacted cytokine protein expression based on the Luminex cytokine profiling assay. Therefore, the EES was shown to modulate mammalian cell gene expression.

**Figure 6.**
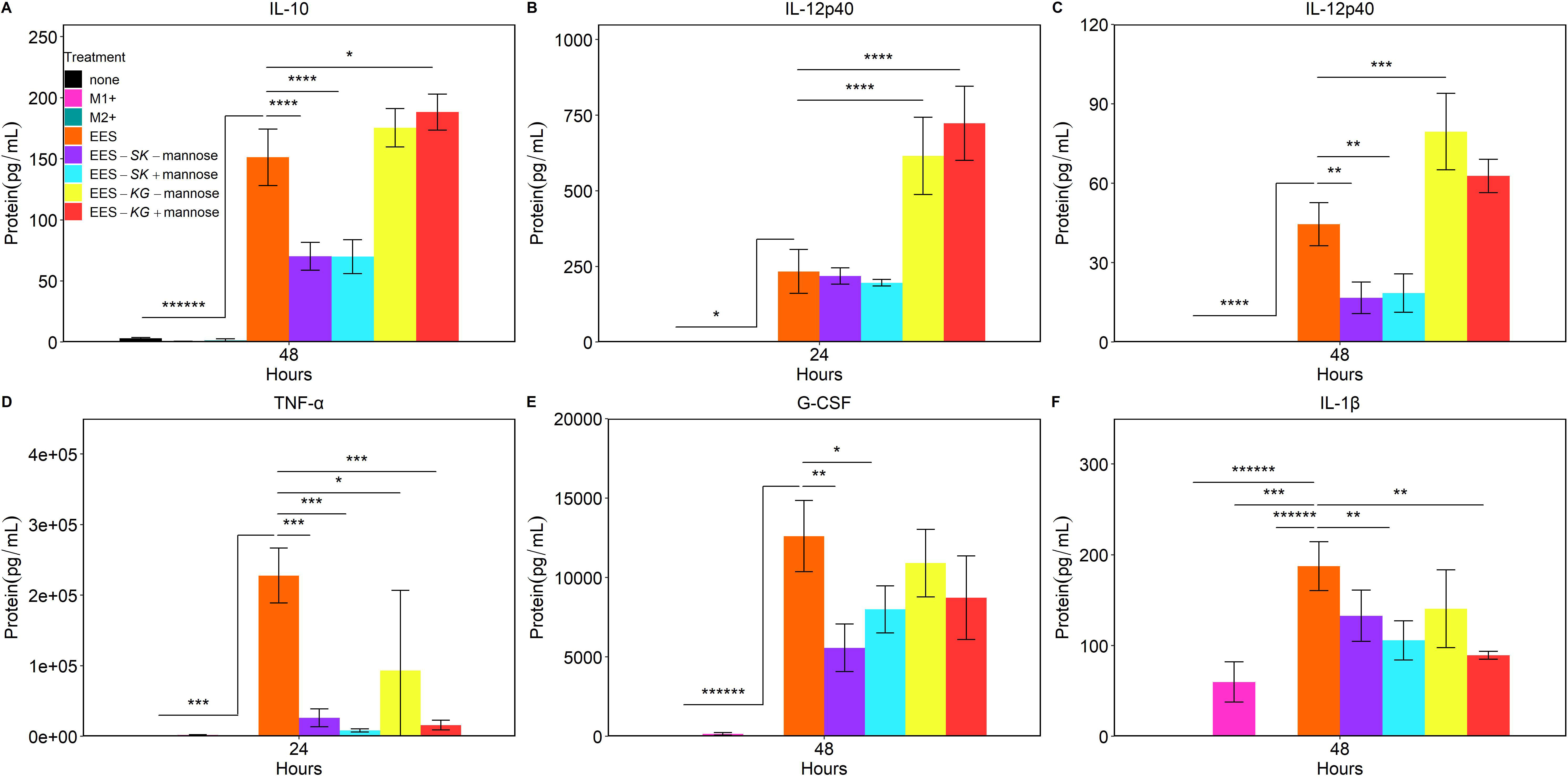
Luminex cytokine profiling assay identifies EES modulation of cytokine expression in J774A.1 cells. Cytokine protein concentration after J774A.1 cells were treated with nothing (none), LPS and IFN-γ (M1+), IL-4 and IL-13 (M2+), EES, EES-*SK* with and without mannose (EES-*SK* -mannose, EES-*SK* +mannose) and EES-*KG* with and without mannose (EES-*KG* -mannose, EES-*KG* +mannose) at 24 and 48 hr post initial treatment. Plotted data is mean ± SD; *p<0.05, **p<0.01, ***p<0.001, ****p<0.0001, ******p<0.000001. Significance shown is comparing EES treatment to all other treatments.

## Discussion

In this study, the EES technology was developed and examined as a tool to alter mammalian cell gene expression. EES viability was identified by replication within the cytoplasm and eventual β-gal protein delivery to the nucleus. Subsequently, the utility of the EES was demonstrated by delivery of transcription factors to the nucleus and the subsequent change in mammalian cell protein expression.

For the EES to successfully impact host cells, a number of events need to occur. First, the EES need to effectively access the cytoplasm of the host cells. Escape of EES from the endosome is essential for survival and function. Induction of the EES to secrete LLO allowed the EES to maintain morphology within the cytoplasm of the host cell (Kortebi et al., 2017) (Figure 2, Movie S1), indicating escape from lysosome-mediated destruction. J774A.1 cells were found to contain an average of 11 EES per cell with 99% of cells containing EES (Table S1). In practice, this percentage, and the EES number per cell, would be varied by manipulating EES according to the application. Second, the EES must remain viable and metabolically active while in the host cell. Initially, this was visualized by EES replication in the cytoplasm. Furthermore, intracellular EES were shown to actively make and deliver β-gal protein to the nucleus. Previous studies also indicated a potential intrinsic NLS associated with *E. coli* β-gal during discovery and testing of modified SV40 signals that caused trafficking to nucleus without the NLS (Kalderon et al., 1984). The mannose-inducible system has been shown to allow transcription even without mannose (Sun and Altenbuchner, 2010), quantification of β-gal in the nucleus also demonstrated some leakiness (Sun and Altenbuchner, 2010). Third, the host cells need to remain viable. Without regulation of EES replication, the fate of a host cell with EES will generally be cell death, due to proliferation of the EES. This was apparent in 10-12% of cells at the final time point (Figure 3A). In this study, the cultures were treated with an increased concentration of gentamicin, above the amount used to kill extracellular bacteria, to kill the intracellular EES, but to further develop the EES chassis, replication must be controlled to optimize the interaction between the EES and host cell. Controlling the EES replication through metabolic control may not be ideal as the EES to produce protein. Genetic control of EES replication machinery could control replication and maintain protein production. In future work, an optimized EES could contain a genetic switch controlling an essential gene responsible for initiation of replication. A key regulator that synchronizes with growth to trigger replication (Witz et al., 2019), such as *dnaA* should be used to regulate genome duplication (Messer, 2002).

Macrophages are plastic as indicated by changes in their cell surface markers and gene expression profiles, which can change quickly as they respond to their environment. In this study this plasticity was modulated by transcription factors produced by EES, and by the EES itself, at 24 and 48 hr after phagocytosis. CD206 has been shown to recognize the surface carbohydrates of pathogens and be triggered by proteases produced by *B. subtilis* (Corvey et al., 2003; García-González et al., 2019; Stahl and Ezekowitz, 1998). Thus, CD206 marker expression is complex and may be increased by nonspecific factors. Additionally, while CD86 provides more clarity because of the marker being regulated directly by inflammatory responses to the EES, the plasticity of the macrophages could cause CD86 expression to change over time (Mantovani et al., 2004). It has been suggested that the products of *B. subtilis* (*i*.*e*. sublancin or exopolysaccharide; EPS) or the treatment of macrophages with *B. subtilis* spores can result in either M1 or M2 activation (Hong et al., 2019; Paynich et al., 2017). While the EES parent *B. subtilis* strain 168 does not produce EPS or spores, a complex reaction to the EES was observed.

Although the changes in cytokine expression vary along the spectrum of macrophage polarization, there are well documented cytokines which are used to identify, broadly, M1 or M2 macrophages. These identifying cytokines are important when studying disease as they provide information on the function and characteristics of the macrophage population. Macrophages shifted towards M1 polarization commonly produce pro-inflammatory cytokines, aiding in destruction of pathogens, including the EES in this case. Alternatively, M2 macrophages decrease inflammation and promote tissue remodeling and repair. These two classifications are helpful but oversimplified; there are complexities in macrophage activation, phenotype and plasticity, as was encountered in this study. In some instances, the EES did act on the host cell as predicted; the upregulation of IL-10 by EES-*KG* and down regulation by EES-*SK* aligns with what is expected of these macrophages (Liao et al., 2011; Rőszer, 2015; VanDeusen et al., 2006). However, in some cases there were results that were not anticipated. Accordingly, the effects of each EES transcription factor need to be considered. For example, KLF4 has been implicated in the reduction of TNF-α (Liao et al., 2011) but did not upregulate G-CSF. The change in IL-12p40 levels is an example of an unexpected result based on known signaling cascades. IL-12p40 is known to be produced during inflammatory phenotypes by nuclear factor kappa light chain enhancer of activated B cells (NF-κB) activating the IL-12p40 promoter (Becker et al., 2001; Fraternale et al., 2013). Therefore, the EES-*KG* strain causing an increase in production of IL-12p40 is unexpected. However, GATA-3 could play a role in IL-12p40 production. The IL-12p40 promoter has a characterized GATA binding site but GATA-3 has not been characterized in terms of impact on this promoter (Becker et al., 2001). Furthermore, there may have been a metabolic impact on cytokine production due to the addition of D-mannose, which impairs glucose metabolism leading to downregulation of IL-1β. Metabolic reprogramming is most likely playing a significant role in response to the chemical inducer of the EES operons and possibly the EES alone (Torretta et al., 2020).

Controlling inflammation as a therapeutic could be an important biomedical application of the EES-*SK* and EES-*KG* strains. For example, inflammation is an important hallmark of cancer, and the presence of immune cells known as tumor associated macrophages (TAM) is often conducive of tumor growth, metastases and poor outcomes (Gharib et al., 2019; Laoui et al., 2011). TAMs are broadly M2 polarized, and IL-10, IL-12p40 and G-CSF all have been shown to play important roles in impacting the tumor microenvironment through regulation of TAMs (Biswas et al., 2006; Kanemaru et al., 2017; Karagiannidis et al., 2020; Laoui et al., 2011) which could be targeted by the EES in cancer bacteriotherapy (Sawant et al., 2020; Yaghoubi et al., 2019, 2020b, 2020a; Zhang et al., 2020). Arthritis represents the other side of immune polarization where homeostasis is driven to a proinflammatory condition (Georgopoulos et al., 1996; Henderson and Pettipher, 1989). Here, modulating macrophages towards the M2 phenotype could reduce inflammation in joints. The EES ability to downregulate TNF-α and upregulate IL-10 shows promise for treating damaging inflammatory conditions such as that present in arthritis (Boehler et al., 2014; Georgopoulos et al., 1996; Henderson and Pettipher, 1989; Iyer and Cheng, 2012; Tran et al., 2015) Manipulation of immune cells, such as macrophages, has been well characterized *in vitro*. However, their use *in vivo* presents problems of low efficacy and lack of innate control (Spiller and Koh, 2017). The use of EES could circumvent these issues since EES can be taken up by phagocytic cells and impact gene expression. Thorough characterization of the impact on primary macrophages and the signaling produced during and after EES delivery and manipulation of those cells will be critical in understanding the impact EES could have on human disease.

Another possible application for the EES could involve delivering clustered regularly interspaced palindromic repeats (CRISPR-Cas9) to mammalian cells as the *Streptococcus thermophilus* CRISPR3-Cas system has been engineered into *B. subtilis* previously to protect against bacteriophage contamination in industrial fermentation (Jakutyte-Giraitiene and Gasiunas, 2016). The functional delivery of protein to the nucleus observed in this study by the EES could facilitate CRISPR-Cas9 gene editing in mammalian cells. EES could be an alternative to nanoparticles, viral methods, cell penetrating peptides and electroporation, especially for *in vivo* delivery and delivery to phagocytic cells (Liu et al., 2017). Additionally, EES could serve as an on-demand production vehicle of CRISPR-Cas9 and could be engineered to generate variations of CRISPR-Cas9 under control of different regulatable systems.

The use of an EES is advantageous when compared to alternative methods of manipulating mammalian cell fates and function. Current methods of manipulating mammalian cell fate include viral vectors, growth factors or signaling molecules (Lienert et al., 2014). Additionally, chimeric antigen receptor T (CAR-T) cells and CRISPR technologies have proven to be potential futures for some therapeutics and are clinically relevant (Larson and Maus, 2021; Lienert et al., 2014; Mollanoori et al., 2018; Salas-Mckee et al., 2019). However, even the more current technologies have limitations for certain applications especially *in vivo* (Larson and Maus, 2021; Lienert et al., 2014; Mollanoori et al., 2018; Salas-Mckee et al., 2019). Also, viral vectors have been shown to be slow as therapeutics within the immune system, specifically in targeting and modulating macrophages, compared to exogenous cytokines (Boehler et al., 2014). The more recent method of manipulating cellular fates using prosthetic networks increases the variety of cargo that can be delivered and provides some control with limitations (Folcher and Fussenegger, 2012; Kemmer et al., 2010; Lienert et al., 2014; Ye et al., 2011). An EES can build on the precedence of prosthetic networks by having the capability to be constructed to generate protein and molecules once in the cytoplasm of mammalian cells for improved control; the method of continuously supplying protein in the form of transcription factors could drastically improve obtaining the desired fate.

Although phagocytic cells present good targets for EES, delivery of EES to different cell types, including normally non-phagocytic cells will be necessary for some future applications. Some bacteria can gain entry to a non-phagocytic host cell after initiating Trigger or Zipper entry processes (Günther and Seyfert, 2018). Protein classes such as internalins or invasins are used by pathogens to gain entry into non-phagocytic cells through well studied mechanisms and could be utilized by the EES (Cossart and Sansonetti, 2004). A few examples of these proteins include internalin A (InlA) and internalin B (InlB) from *L. monocytogenes* which bind to E-cadherin and tyrosine kinase Met, respectively, or surface cell antigen 2 protein (Sca2) from *Rickettsia conorii* to invade via actin (Chao et al., 2021; Cossart and Sansonetti, 2004). Many of these bacteria are then contained within a membrane-bound vacuole, resulting in death, or survival after escape into the cytoplasm (Cossart and Sansonetti, 2004). The above processes can be adapted into an EES with a phospholipase/hemolysin component. LLO, as observed in this study, may be necessary for escape even after use of an internalin or invasin. These virulence factors, like LLO, would need to be properly regulated. Once *in vivo*, tools for targeted delivery and increasing tropism of the EES will be necessary to improve potential therapeutic applications of the EES.

Studies of pathogens and their virulence factors could be used to inform future development of the EES. The EES will utilize this deep characterization for defined control within the host cell cytoplasm and their interaction with nuclear processes. This study demonstrated the utility of EES to direct cell gene expression and alter the fates of mammalian cells. This use of the EES as a tool to change mammalian cell gene expression only begins to demonstrate the possibilities, and the impact of EES control modules in mammalian cells could have a significant impact on treatment of a wide variety of diseases by altering the fates and functions of cells in the body. *B. subtilis* serves as an ideal chassis for development and optimization of the EES capable of existing viably in the cytoplasm and delivering protein to the nucleus of a mammalian cell to alter cellular fates.

### Limitations of this study

Using only the J774A.1 cell line potentially limits the scope of this study. Future work will pursue characterizing EES impact on primary macrophages to bolster translatability.

## Supporting information

Key resource table

Oligo list

## Acknowledgements

The authors would like to thank Dr. Daniel A. Portnoy for the *B. subtilis* LLO strain, Dr. Jens Schmidt for advice on imaging and Dr. Lee Kroos for *B. subtilis* plasmid pDR111. The authors would also like to thank Dr. Melinda Frame of the Michigan State University Center for Advanced Microscopy for confocal imaging, Dr. Matthew Bernard of the Michigan State University Flow Cytometry Core for flow cytometry aid and Luminex cytokine profiling analysis. The authors would like to acknowledge Chima V. Maduka for his thoughtful and thorough review of the manuscript and providing crucial insights. The authors would like to acknowledge the James and Kathleen Cornelius Endowment for financially supporting the project. The graphical abstract figure along with Figure 1 and S1 were created using BioRender.com.

## Author contributions

Cody S. Madsen conceptualized *Bacillus subtilis* as a chassis organism, developed all the EES constructs as the platform technology, jointly developed and performed all experiments, jointly developed and analyzed all data to make figures and was one of the primary authors of the manuscript. Dr. Ashley V. Makela jointly developed and performed all experiments, jointly developed and analyzed all data to make figures and was one of the primary authors of the manuscript. Emily M. Greeson significantly contributed to the development of the EES platform technology, jointly developed and significantly contributed to the writing of this manuscript. Dr. Jonathan W. Hardy acquired the *B. subtilis* LLO strain from Dr. Daniel A. Portnoy, significantly contributed to the development of the EES platform technology and significantly contributed as an author to this manuscript. Dr. Christopher H. Contag conceptualized the initial concept of the EES, supervised the studies, contributed to the experimental design, provided the resources, reviewed data and edited the manuscript.

## STAR★Methods

### Data and code availability

All raw data, EES constructs and R scripts will be made available upon request by the corresponding author. Plasmids used to produce EES constructs will be submitted to Addgene after manuscript publication. All R scripts were written with established packages.

### *B. subtilis* LLO constructs

Constructs were inserted into the genome of *B. subtilis* LLO (**EES**) at the *amyE* locus using a homologous recombination plasmid (pDR111 (Rokop et al., 2004), a gift from Dr. Lee Kroos). The pDR111 plasmid was transformed into *B. subtilis* using a natural competence protocol and constructs were selected for by spectinomycin then confirmed by PCR amplification out of the genome (Harwood and Cutting, 1990). *B. subtilis* expressing IPTG-inducible LLO was provided by Dr. Daniel Portnoy. The constructs include the *lacZ, Stat-1Klf6* and *Klf4Gata-3* genetic cassettes (Supplemental item 1). *B. subtilis* LLO was designed to secrete β-gal to the nucleus through the twin-arginine translocation (Tat) pathway by synthesizing the PhoD signal peptide (Tjalsma et al., 2000) and the amino acids from 126-132 of the simian virus (SV) 40 nuclear localization signal (NLS) (Kalderon et al., 1984) together and connecting both to *lacZ* when inserting into pDR111 using Gibson assembly after *lacZ* was amplified from the pST5832 plasmid (Addgene). The same construct was engineered without the SV40 signal to confirm specific delivery to the nucleus. The *lacZ* gene and the synthesized PhoD signal peptide plus SV40 NLS were cloned into the NheI restriction site in pDR111 using Gibson assembly. Initially, the β-gal secretion strain was controlled by an IPTG-inducible promoter (Phyper-spank). However, previous studies have shown the IPTG system is limited in controlling protein production so these constructs were engineered to be controlled by the mannose-inducible system amplified from *B. subtilis* ZB307 strain genome (Sun and Altenbuchner, 2010). The mannose promoter and regulator were cloned into the pDR111 plasmid to replace the Phyper-spank promoter and LacI regulator using Gibson assembly. Accordingly, the *lacZ* gene with the same design was cloned in the NheI restriction site present in the pDR111 mannose plasmid using Gibson assembly. The *Stat-*1 and *Klf6* genes were synthesized by IDT as a custom gene and Gblock respectively from the coding sequences obtained from Uniprot. The *Stat-1Klf6* operon was fused by ligation at an introduced EagI restriction site between the genes during cloning into the pDR111 mannose plasmid by restriction cloning into the SalI and NheI restriction sites (Figure S7). The *Klf4* and *Gata-3* genes were synthesized as Gblocks from IDT and fused using the same method as the *Stat-1Klf6* operon. The *Klf4Gata-3* operon was cloned into the pDR111 mannose plasmid using restriction cloning at the SalI and SbfI restriction sites after the SbfI cut site was introduced into the multiple cloning site by inverse PCR then digesting both ends with SbfI and re-ligating the pDR111 mannose plasmid (Figure S7). All constructs were confirmed by restriction digest, sequencing and functionality tests.

### EES culture growth conditions

The EES was grown under the same conditions for all experiments. Each EES construct was grown in Luria-Bertani Miller broth (LB) with the appropriate antibiotic. *B. subtilis* LLO was grown in LB with chloramphenicol (10 µg/mL) and all constructs that were integrated into the *amyE* were grown with spectinomycin (100 µg/mL). The overnight cultures were grown for 16 hr at 37°C and 250 RPM. All constructs were integrated into the genome of the EES which allowed for expression of constructs without antibiotics during co-incubation with J774A.1 cells.

### EES delivery protocol

The following conditions were utilized to induce EES delivery, unless otherwise described. J774A.1 monocyte/macrophage cells (ATCC, VA, USA) were maintained at 37°C and 5% CO_2_ in DMEM (ThermoFisher, MA, USA), supplemented with 10% fetal bovine serum (FBS). Cells were tested negative for mycoplasma using the MycoAlert PLUS Mycoplasma Detection Kit (Lonza, xx). Once cells were confluent, they were seeded onto a plate or 4-well chambered imaging slide and allowed to adhere overnight (described below) when an estimation of total number of cells was made based on confluency. EES was added at an optimized MOI of 25:1 for all experiments besides testing of host cell viability, along with IPTG (500 µM) to induce expression of LLO with or without protein of interest. EES and J774A.1 cells were then co-incubated at 37°C and 5% CO_2_ for 1 hr. J774A.1 cells were then washed three times with PBS and new medium was added containing gentamicin (4 µM) to eliminate any remaining extracellular EES. Co-incubation continued for 3 hr at 37°C and 5% CO_2_ prior to imaging or preparation for microscopy (described below) (Figure S1).

### EES antibody staining for phagosomal escape

After fixation with 4% PFA, cells were permeabilized using 0.3% Triton X-100 (ThermoFisher) followed by a blocking step containing 0.3% Triton X-100 and 5% normal goat serum (ThermoFisher, cat# 31872,). The EES location was determined by incubating a rabbit anti-subtilisin antibody (1:50, Antibodies-online, PA, USA, cat# ABIN958907) at 4°C overnight followed by a goat anti-rabbit IgG Dylight 650 (1:4000, Novus, CO, USA, cat# NBP1-76058) secondary antibody at room temperature (RT) for 2 hr. Phagosome formation or destruction was shown by incubating an anti-Lamp-1 (Kortebi et al., 2017) antibody (1:100, AbCam, MA, USA, cat# ab25245) at 4°C overnight followed by a goat anti-rat IgG Alexa Fluor 555 (1:1000, Invitrogen, CA, USA, cat# A-21434) secondary antibody at RT for 2 hr. Nuclei were counterstained by incubating cells with Hoechst 33342 (1 µg/mL) for 10 minutes (min) at RT. Membranes of the cells were stained by incubating cells with PKH67 green-fluorescent cell linker kit (10 µM, Sigma MO, USA,) for 10 min at room temperature. Slides were then coverslipped using Fluoromount-D mounting media (Southern Biotech, AL, USA). Slides were imaged using confocal microscopy (described below).

### Confocal imaging

Confocal microscopy was performed using a Nikon A1 CLSM (Nikon, NY, USA) microscope to determine EES escape from the phagosome complex by imaging J774A.1 cells that had been treated with the EES with and without IPTG. Imaging was performed using a 60x oil objective and 1.5x zoom and using filter sets for DAPI (Hoechst 33342), GFP (PKH67), TRITC (Alexa Fluor 555 for LAMP-1) and Cy5 (Dylight 650 for *B. subtilis*). Z-stacks were taken at 0.5 µm steps to confirm location of EES within host cells. Images were analyzed using NIS-Elements AR Software (Nikon) and background noise was reduced by using Nikon denoise.ai algorithm. The 3-dimensional volume images and cutaways were produced by the Alpha display mode. The Alpha display mode was also used to generate 3-dimensional videos to display z-depth location of the EES during escape from the phagosomes. The data presented herein were obtained using instrumentation in the MSU Flow Cytometry Core Facility. The facility is funded in part through the financial support of Michigan State University’s Office of Research & Innovation, College of Osteopathic Medicine, and College of Human Medicine.

### J774A.1 viability by MTS assay and Flow cytometry

The impact of the EES on J774A.1 cell viability was determined by using the EES delivery protocol then performing analysis using an MTS assay and flow cytometry throughout the time (0, 1 and 4 hr) of interaction between the EES and host cells at different MOIs (10:1, 25:1, 50:1), no IPTG induction and no treatment, with biological triplicates (n = 3) for each time and condition in MTS assay. Flow cytometry was performed with a single biological replicate (n = 1) to compare to the MTS assay for same general trend. At each time point for the MTS assay, J774A.1 cells were washed once with PBS then MTS reagent (Abcam) was added at a 10-fold dilution with DMEM then added to cells. After addition, the reagent was incubated with cells for 30 min at 37°C then absorbance was read at 490 nm. All treatment conditions were compared to J774A.1 cells alone to elucidate any differences in loss of viability of the J774A.1 cells due to the treatment conditions. Standard one-way ANOVA with Tukey post-hoc test was used to determine statistically different values in MTS assay. Flow cytometry was performed with a single biological replicate (n = 1) to compare to MTS assay for same general trend. Cells were analyzed at the 4 hr time point, with different MOIs (10:1, 25:1, 50:1), no IPTG induction and no treatment. Cells were washed once with 1X PBS followed by trypsinization to detach the cells. Cells were collected, washed once with 1X PBS and incubated with 1:750 Zombie NIR viability dye in 100 µl 1X PBS for 20 min, room temperature, in the dark. Cells were centrifuged, washed twice and resuspended in 100 µl flow buffer.

### Live Cell Imaging

EES was internalized into J774A.1 cells as described above, using a 96-well black glass-bottom plate (40,000 cells/well; Greiner Bio-One, Austria, cat# 655892). EES were pre-stained using a TRITC fluorescent dye, CellTracker Orange CMRA Dye (Invitrogen, C34564). The EES was spun down (10,000 x g) for 2 min then resuspended in CellTracker Orange CMRA Dye (2 µM) in PBS then incubated at 37°C and 250 RPM for 25 min. Afterwards, the EES were centrifuged (10,000 x g) and washed three times before adding to J774A.1 cells. Live cell imaging was performed on a Leica DMi8 Thunder microscope equipped with a DFC9000 GTC sCMOS camera and LAS-X software (Leica, Wetzlar, Germany). Cells were maintained at 37°C and 5% CO_2_ in Fluorobrite medium during the imaging session. Fluorescent images were acquired using a TRITC filter set. Brightfield and fluorescent images were acquired consecutively, using a 63x oil objective every 1 hr beginning at 1 hr post co-incubation and continuing until 4 hr post incubation Z-stacks were taken at all time points at 0.4 µm steps to confirm EES presence within cytoplasm. EES presence in J774A.1 cells were quantified using Fiji (ImageJ) software and cell counter plugin. An area of 2090 µm by 1254 µm in each well was imaged and used to perform this quantification.

### EES β-gal protein secretion

EES engineered to secrete β-gal (EES-*lacZ*) with and without an NLS (EES*-lacZ-*NLS and EES*-lacZ-*no NLS) were allowed to internalize into J774A.1 cells as described above, using 4-well chambered slides (75,000 cells/well, ThermoFisher, cat #154917). Incubation was also performed using EES-*lacZ*-no NLS and the supernatant from an induced EES*-lacZ-*NLS culture. The second incubation step, after adding gentamicin (4 µg/mL) and with/without D-mannose (1% w/v) for the β-gal secretions strains, was performed for 3 hr. During extended trafficking studies, the cells were washed gently 3 times with 1X PBS then a high concentration of gentamicin (25 µg/mL) was added after 3 hr then incubation continued for an additional 21 hr. The cells were then fixed with 4% paraformaldehyde (PFA) for 10 min prior to preparation for microscopy (described below).

### β-gal antibody staining

After fixation with 4% PFA, cells were permeabilized using 0.3% Triton X-100 (ThermoFisher) followed by a blocking step containing 0.3% Triton X-100 and 5% normal goat serum (ThermoFisher). β-gal secretion to the nucleus was identified using fluorescence imaging after incubation with an anti-β-galactosidase (*E. coli*) antibody-rabbit (1:100, Biorad, CA, USA, cat# AHP1292GA) at 4°C overnight followed by a goat anti-rabbit IgG Dylight 650 (1:3000, Novus) secondary antibody at RT for 2 hr. Nuclei were counterstained by incubating cells with Hoechst 33342 (1 µg/mL) for 10 min at room temperature. Slides were then coverslipped using Fluoromount-D mounting media (Southern Biotech, AL, USA). Epi-fluorescent microscopy was performed using a Nikon Eclipse Ci-L microscope equipped with a CoolSNAP DYNO camera for fluorescent imaging and NIS elements BR 5.21.02 software (Nikon). Images were acquired with a 40x phase contrast objective and for fluorescent imaging DAPI and Cy5 filter sets were used. Nuclear SNR was quantified by imaging in random areas in each corner of the well and in the center of the well. At least 5 random images totaling an area of 1085 µm by 825 µm were utilized in drawing regions of interest (ROIs) around nuclei of J774A.1 cells to quantify Cy5 fluorescence by utilizing Hoechst 33342 counterstain to determine nuclei location. Nuclear SNR was calculated by using mean fluorescence intensity of ROI region around nuclei of J774A.1 cells in each of the five random images (n = 50 random individuals) divided by standard deviation of noise in the well. Standard deviation of noise in well was determined by drawing 3 ROIs (550 µm^2^) in background areas of each image then calculating standard deviation of the mean fluorescence from all of the ROIs drawn in background areas. Statistics was determined using Brown-Forsythe and Welch ANOVA with Dunnett T3 post-hoc test and all treatment conditions were compared.

### EES-modulation of J774A.1 cell gene expression

EES containing mannose-regulatable *Stat-1* and *Klf6* (EES-*SKl*) plus *Klf4* and *Gata-3* transcription factors (EES-*KG*) were internalized into J774A.1 cells as described above (EES β-gal protein secretion), using a 6-well plate (Corning Costar #3516). These transcription factors were secreted for 3 hr then allowed to be trafficked for an additional 21 hr as described above (EES β-gal protein secretion). Additionally, for flow cytometry and Luminex characterization, media was changed at 24 hr and incubation continued for an additional 24 hr. For flow cytometry, Accutase (Sigma, cat# A6964) was added to J774A.1 cells exposed to all treatments at both 24 and 48 hr for 30 min, and then cells were removed for analysis (described below). For Luminex cytokine profiling (Millipore Sigma, MA, USA), the supernatant from J774A.1 cells exposed to all conditions was removed at both 24 and 48 hr and then analysis was performed to quantify cytokines produced (described below). Non-stimulated J774A.1 cells were assumed to be at resting state (M0). J774A.1 cells were polarized with interferon-gamma (IFN-γ) and lipopolysaccharide (LPS) (M1, 100 ng/ml each) or interleukin (IL)-4 and/or IL-13 (M2, 100 ng/ml each) to be used as positive controls. Furthermore, J774A.1 cells were treated with the EES, EES-*SK* and EES-*KG*. The EES with operons were treated with and without mannose as described in EES β-gal secretion and EES alone was not as no difference in impact on J774A.1 cells was observed in previous flow experiments. All treatment conditions were performed in biological triplicates (n = 3).

### Flow cytometry

Samples were prepared for staining by resuspending 1×10^6^ cells in 100 µl 1X PBS in a 96-well round bottom plate. Samples were first incubated with Zombie NIR viability dye (1:750, Biolegend, San Diego, CA, USA; Cat# 423105) for 15 min at RT in the dark. Cells were washed once with flow buffer, followed by incubation with TruStain FcX™ PLUS (anti-mouse CD16/32) Antibody (Biolegend, Cat#156603; 1.25 µl/sample) for 10 min on ice. Alexa Fluor® 647 anti-mouse CD86 Antibody (2.5 ul/sample; Biolegend; Cat#105020) was then added and incubated for 20 min at RT in the dark. Cells were washed twice with flow staining buffer and fixed with 4% paraformaldehyde for 10 min in the dark. Cells were permeabilized (0.3% TritonX-100 in flow wash buffer) followed by incubated with Brilliant Violet 421™ anti-mouse CD206 (MMR) Antibody (1.25 µl/sample; Biolegend; Cat# 141717) for 20 min at RT in the dark. Cells were washed twice with flow buffer and resuspended in a final volume of 100 µl for flow cytometry analysis using the Cytek Aurora spectral flow cytometer (Cytek Biosciences, CA, USA). Fluorescence minus one controls were used to assess fluorescent spread and for gating strategies. Flow cytometry data was analyzed with the software FCSExpress (DeNovo Software, CA, USA). A standard one-way ANOVA was used to determine statistically different MFI values amongst all groups. A full fit two-way mixed model ANOVA was used to determine statistical changes in CD86+/CD206+ population values amongst cells containing and not containing an EES, and between EES treatment conditions. Multiple comparisons were made using a Tukey post-hoc test. The data presented herein were obtained using instrumentation in the MSU Flow Cytometry Core Facility. The facility is funded in part through the financial support of Michigan State University’s Office of Research & Innovation, College of Osteopathic Medicine, and College of Human Medicine.

### Luminex cytokine profiling assay

Cell culture supernatant from the above flow cytometry experiment were stored at -20°C until ready for use. Supernatant was analyzed for CCL2 (MCP-1), CCL3 (MIP-1a), CCL11 (Eotaxin), CXCL2/MIP-2, G-CSF, IL-1α, IL-1β, IL-4, IL-6, IL-10, IL-12p40, IL-12p70, IL-13, INFγ, LIF, TNF-α and VEGFα cytokine expression. Cytokine levels of cell supernatants were measured using a MCYTOMAG-70K-17 Mouse Cytokine Magnetic Multiplex Assay (Millipore Sigma) using a Luminex 200 analyzer instrument (Luminex Corp, USA) according to the manufacturer’s instructions. Standard one-way ANOVA with Tukey post-hoc test was used to determine statistically different values amongst all treatment groups.

### ICC confirming EES manufacturing and delivering transcription factors

Protocol for EES modulation of J774A.1 cell gene expression was used as the template protocol for this experiment. The experiment was performed using a 96-well black glass-bottom plate (40,000 cells/well; Greiner Bio-One). However, J774A.1 cells were fixed at 3 hr post mannose addition to reveal EES delivering transcription factors to host cells using antibodies against the transcription factor of interest. Each transcription factor was stained for with the appropriate antibody individually within the wells. The EES was stained prior to addition to host cells using Invitrogen CellTracker Orange CMRA Dye (Invitrogen) as in live cell imaging. After fixing, cells were permeabilized using 0.3% Triton X-100 (ThermoFisher) followed by a blocking step containing 0.3% Triton X-100 and 5% normal goat serum (ThermoFisher). Cells were then incubated with rabbit anti-*Stat-1* (1:50, MyBiosource, CA, USA, cat# MBS125754), rabbit anti-*Klf6* (1:50, MyBiosource, cat# MBS8307089), rabbit anti-*Klf4* (1:20, MyBiosource, cat# MBS2014661) or rabbit anti-*Gata-3* (1:100, MyBiosource, cat# MBS8204267) at 4°C overnight followed by a goat anti-rabbit IgG Dylight 650 (1:3000, Novus) secondary antibody at RT for 2 hr. Nuclei were stained by incubating cells with Hoechst 33342 (1 µg/mL) for 10 min at room temperature. Membranes of the cells were stained by incubating cells with PKH67 green-fluorescent cell linker kit (10 µM) for 10 min at room temperature. Imaging was performed using Leica DMi8 Thunder microscope equipped with Leica DFC9000 GTC sCMOS camera and Leica LAS-X software (Leica). DAPI (Hoechst 33342), FITC (PKH67), TRITC (CellTracker Orange CMRA Dye) and Cy5 (Dylight 650 for transcription factors) filters were used and imaged using a 40X objective to confirm EES delivery of transcription factors.

### Statistical analysis

Statistical analyses were performed using Prism software (9.2.0, GraphPad Inc., La Jolla, CA). Statistical tests are identified for each method. Data are expressed as mean +/- standard deviation; *p*<.05 was considered a significant finding. Plotting was performed using R version 4.0.4 with the following packages: ggplot2, dplyr, reshape2, ggsignif and ggpubr.

## Title for videos

**Movie S1. Cutaway of z-depth using all fluorescent channels to reveal EES location within J774A.1 cells with addition of IPTG and related to Figure 2**

**Movie S2. Cutaway of z-depth using all fluorescent channels to reveal minimal EES escape or destruction in phagosomes of J774A.1 cells when IPTG is not added and related to Figure 2**

