## Supplementary material for "Engineered endosymbionts capable of directing mammalian cell gene expression": Key resource table

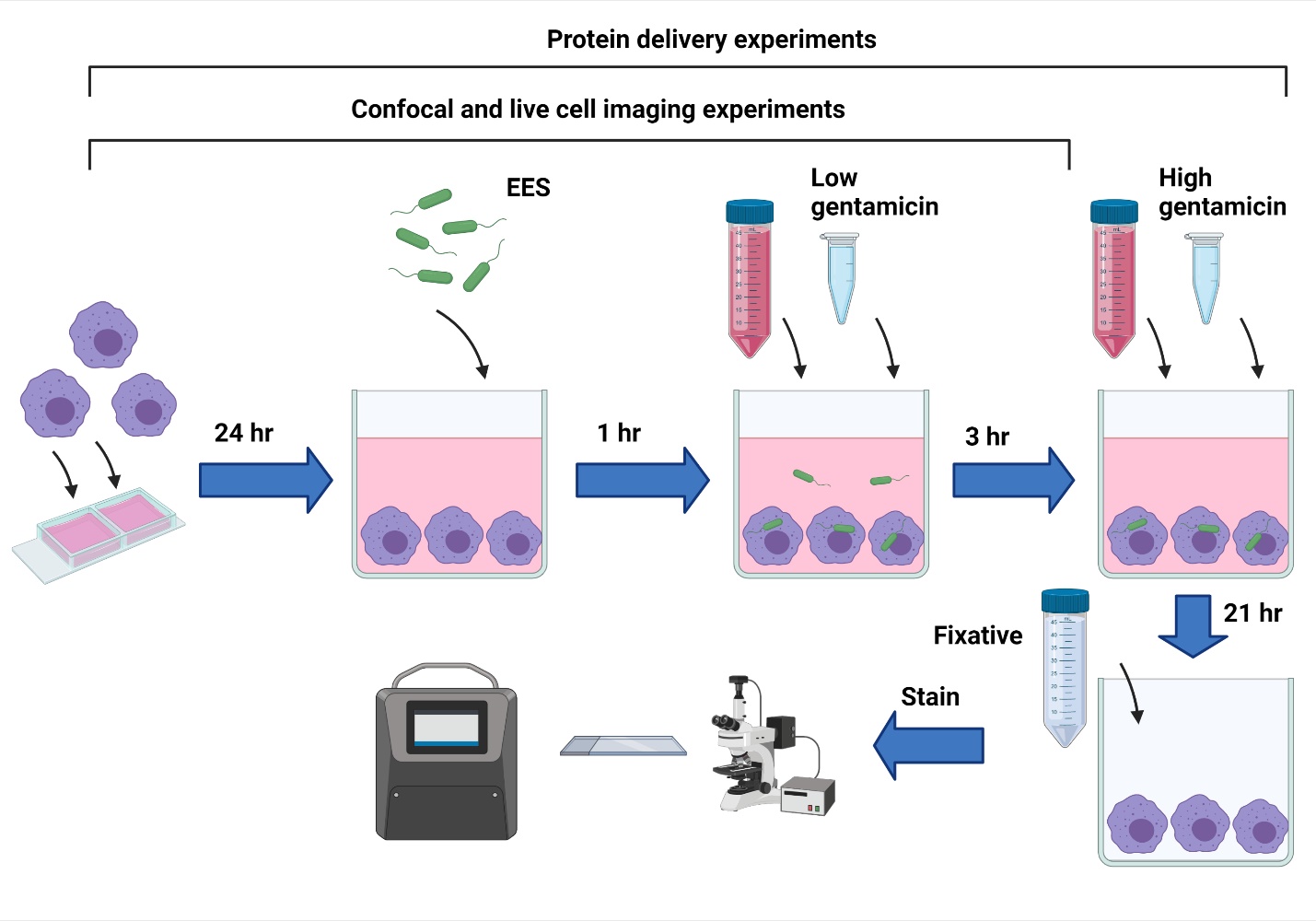

Figure S1: Diagram of method to deliver EES and analyze interaction with host cells and related to Figure 1

General approach for co-incubating EES with host J774A.1 cells and timeline for the interaction. EES are allowed to incubate with J774A.1 cells for 1 hr with the appropriate inducer depending on condition before a low concentration of gentamicin is added to eliminate extracellular EES. Incubation continues for 3 more hours with a second inducer before a high concentration of gentamicin is used to eliminate intracellular EES. Incubation is continued for an extended period of time to determine impact on host cells by imaging or other methods such as cytokine profiling.

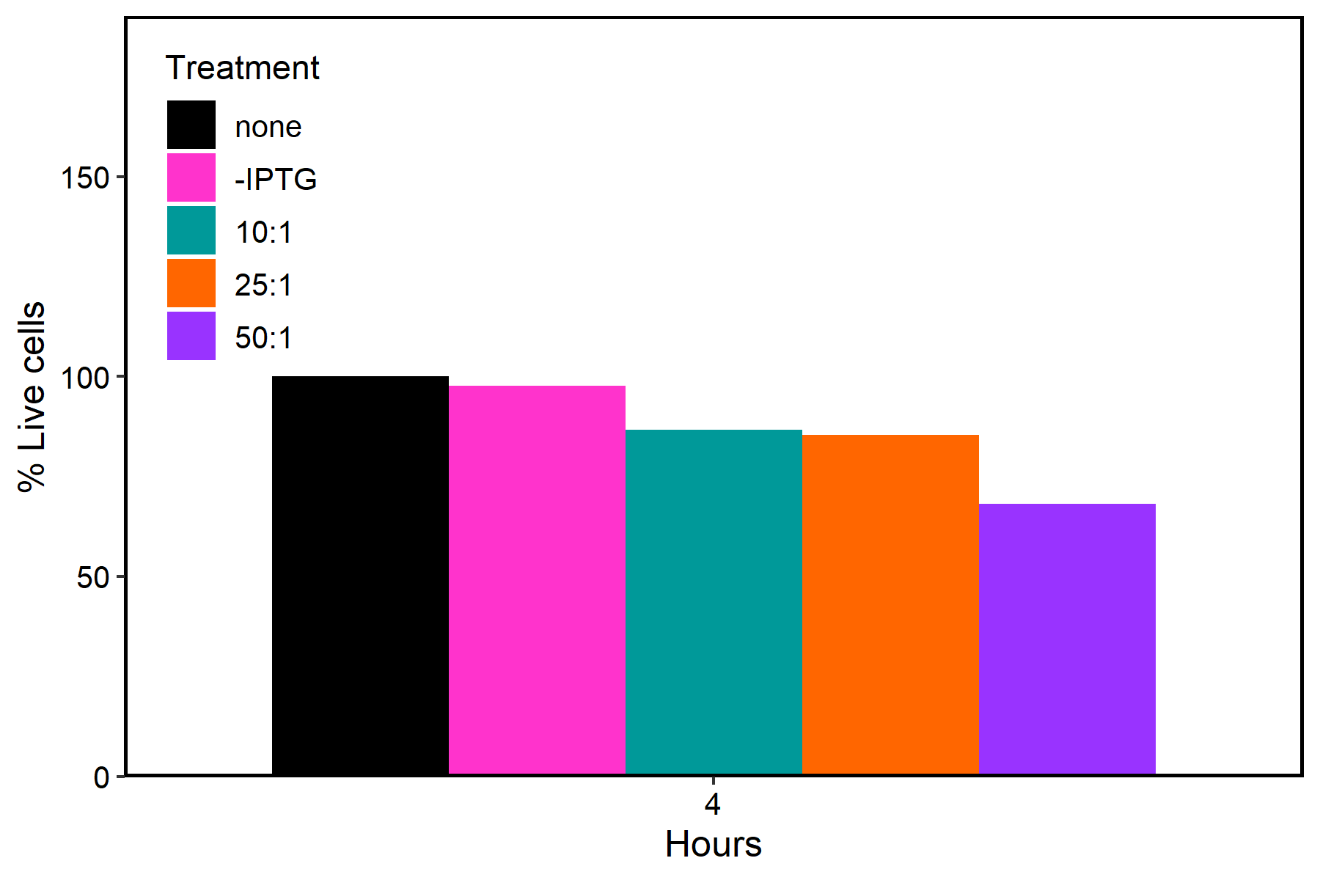

Figure S2: Flow cytometry analysis of viable J774A.1 cells and related to Figure 3A.

J774A.1 cells were treated with EES exposed to IPTG at different MOIs and to EES without IPTG at a 25:1 MOI for 4 hr then removed from the wells. Host cells were then stained with a viability dye and analyzed using flow cytometry. Experiment was performed with one biological replicate (n = 1) to test for the same trend as the MTS assay. Number of events>20,000 cells.

Table S1: Quantification table of EES interaction with J774A.1 cells and related to Figure 3B.

Quantification of EES presence within J774A.1 cells using a 25:1 MOI without IPTG, and different MOI with IPTG at 1 and 2 hr post EES addition.

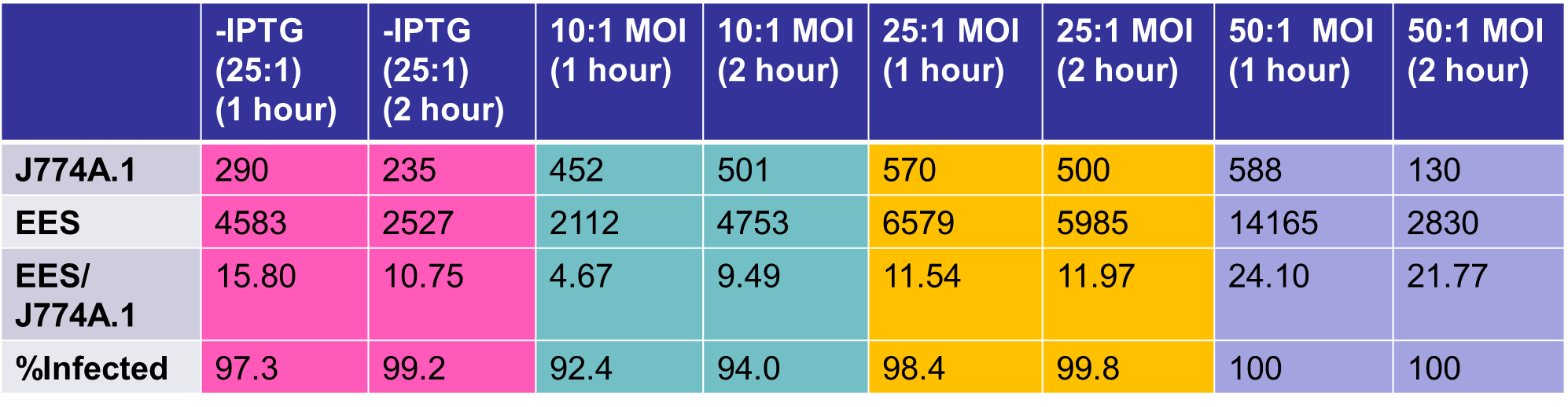

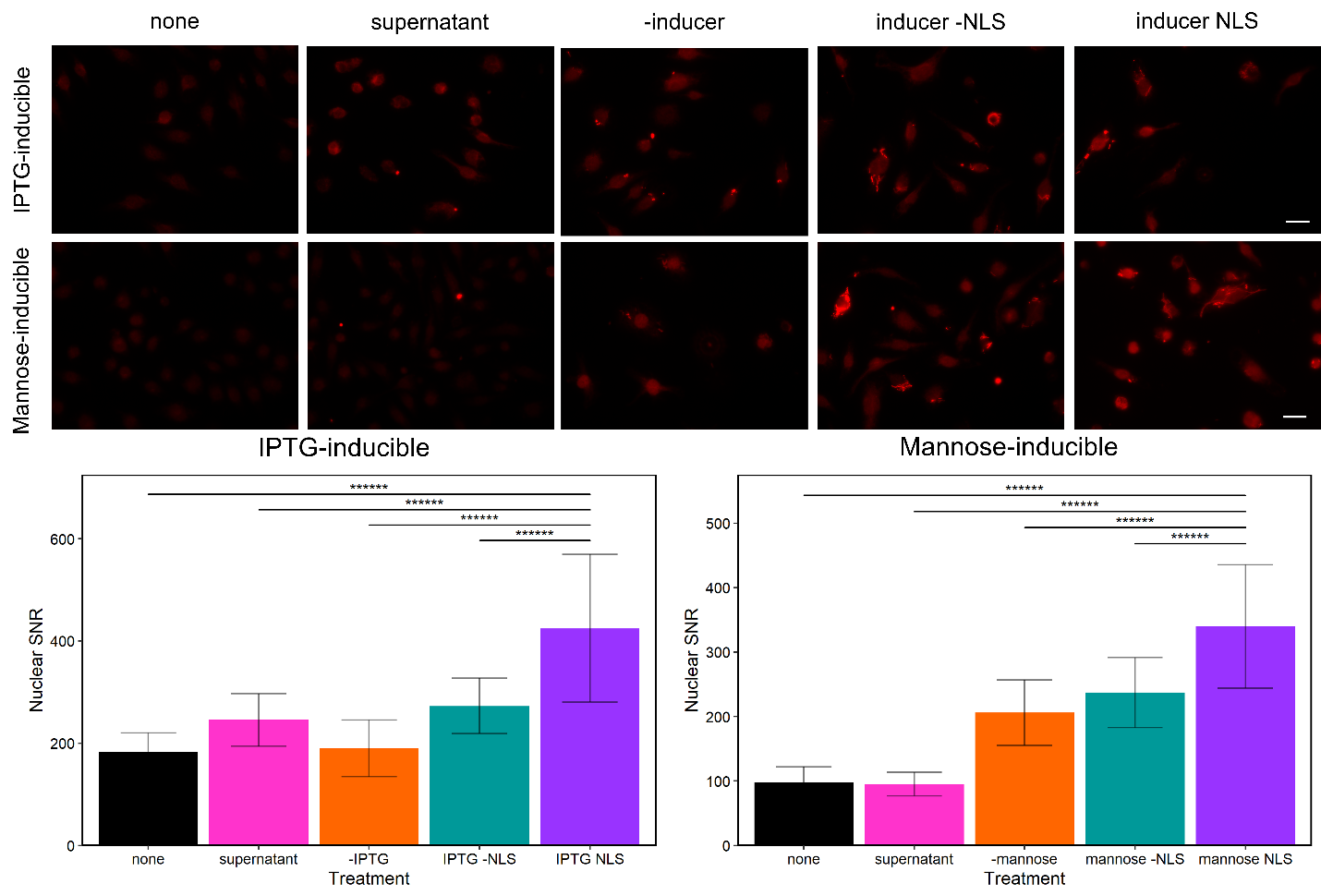

Figure S3. Intracellular EES-β-gal secretes β-gal to nuclei of J774A.1 cell using IPTG- and mannose-inducible systems and related to Figure 4

Fluorescence of nuclei in J774A.1 cells with no EES (none), J774A.1 cells incubated with β-gal collected as supernatant from induced EES-lacZ-NLS (supernatant), J774A.1 cells incubated with uninduced EES-lacZ-NLS (-inducer), J774A.1 cells incubated with induced EES-lacZ-no NLS (inducer -NLS) and J774A.1 cells incubated with induced EES-lacZ-NLS (inducer NLS). Data is mean ± SD; ******p<0.000001. Scale bars = 20 µm.

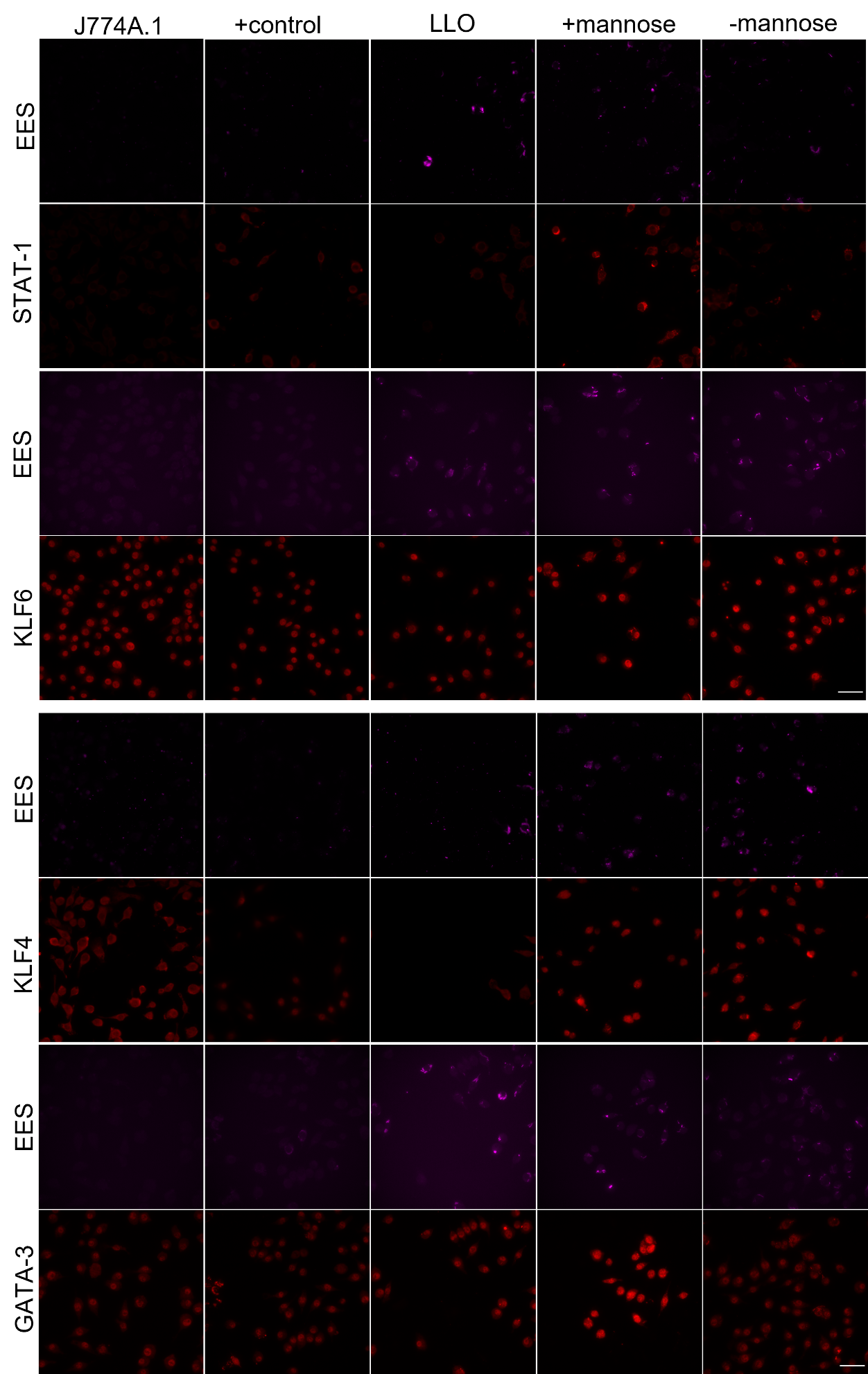

Figure S4. Intracellular EES production and secretion of mammalian transcription factors

Fluorescence microscopy identifies EES (magenta) and transcription factors (red) in J774A.1 cells treated with nothing (J774A.1), positive control (STAT-1/KLF6: LPS and IFN-γ or KLF4/GATA-3: IL-4 and IL-13), EES-LLO (LLO), EES-*SK* or EES-*KG* not induced (-mannose) or induced (+mannose). Each row dynamic range was scaled to the image with the highest fluorescence intensity to compare conditions. Scale bars = 50 µm.

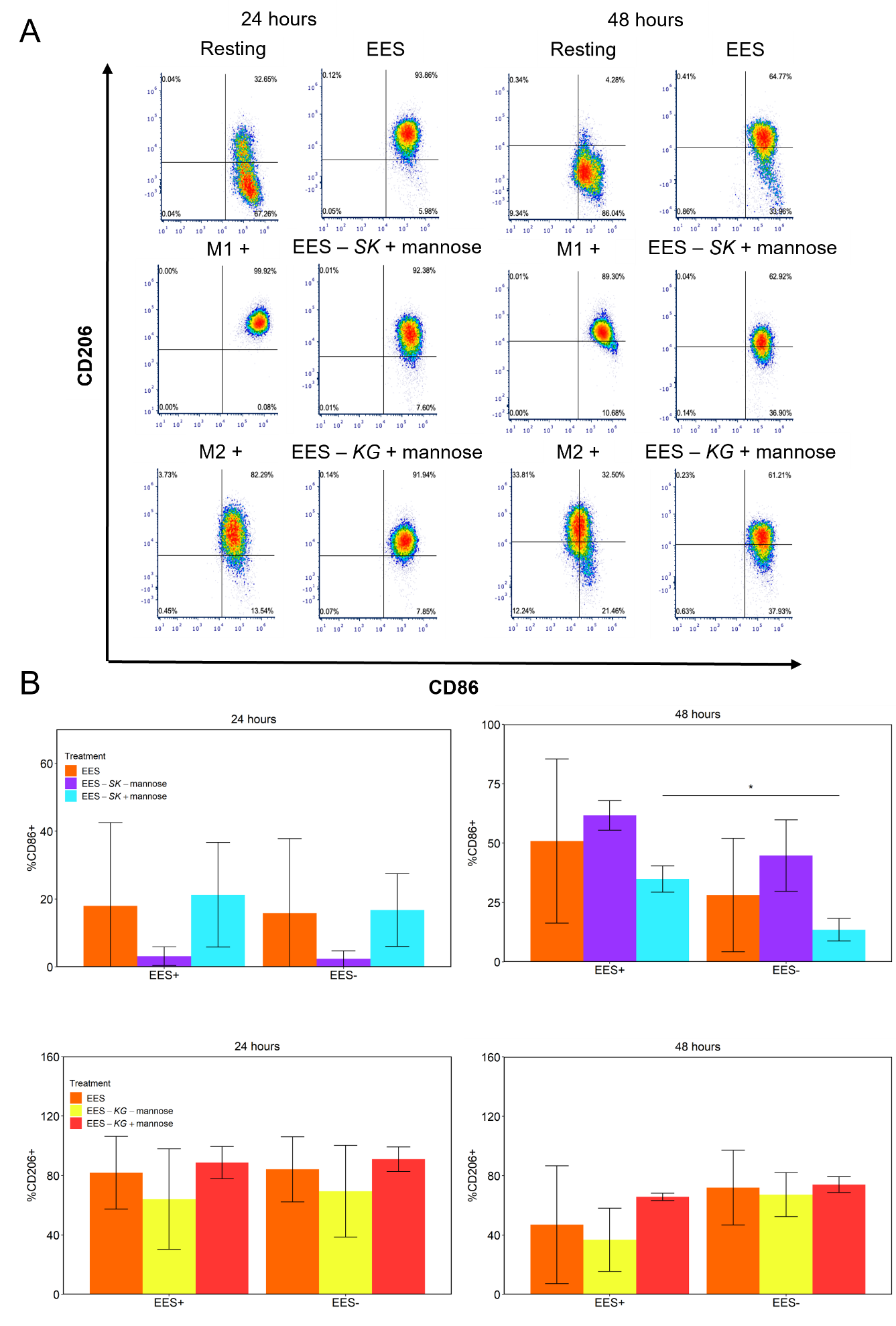

**Figure S5. Further flow cytometry analysis revealing impact of EES on J774A.1 cell gene expression and related to Figure 5**

Flow density plots representative of the histograms display cell population gene expression (A). The populations were analyzed to reveal percentage of CD86 and CD206 expressing cells then separated based on presence of EES stain (B). J774A.1 cells were treated with nothing (none), LPS and IFN-γ (M1+), IL-4 and IL-13 (M2+), EES, EES-*SK* with and without mannose (EES-*SK* -mannose, EES-*SK­* +mannose) and EES-*KG* with and without mannose (EES-*KG* -mannose, EES-*KG* +mannose) at 24 and 48 hr post initial treatment. Plotted data is mean ± SD; **p*<0.05.

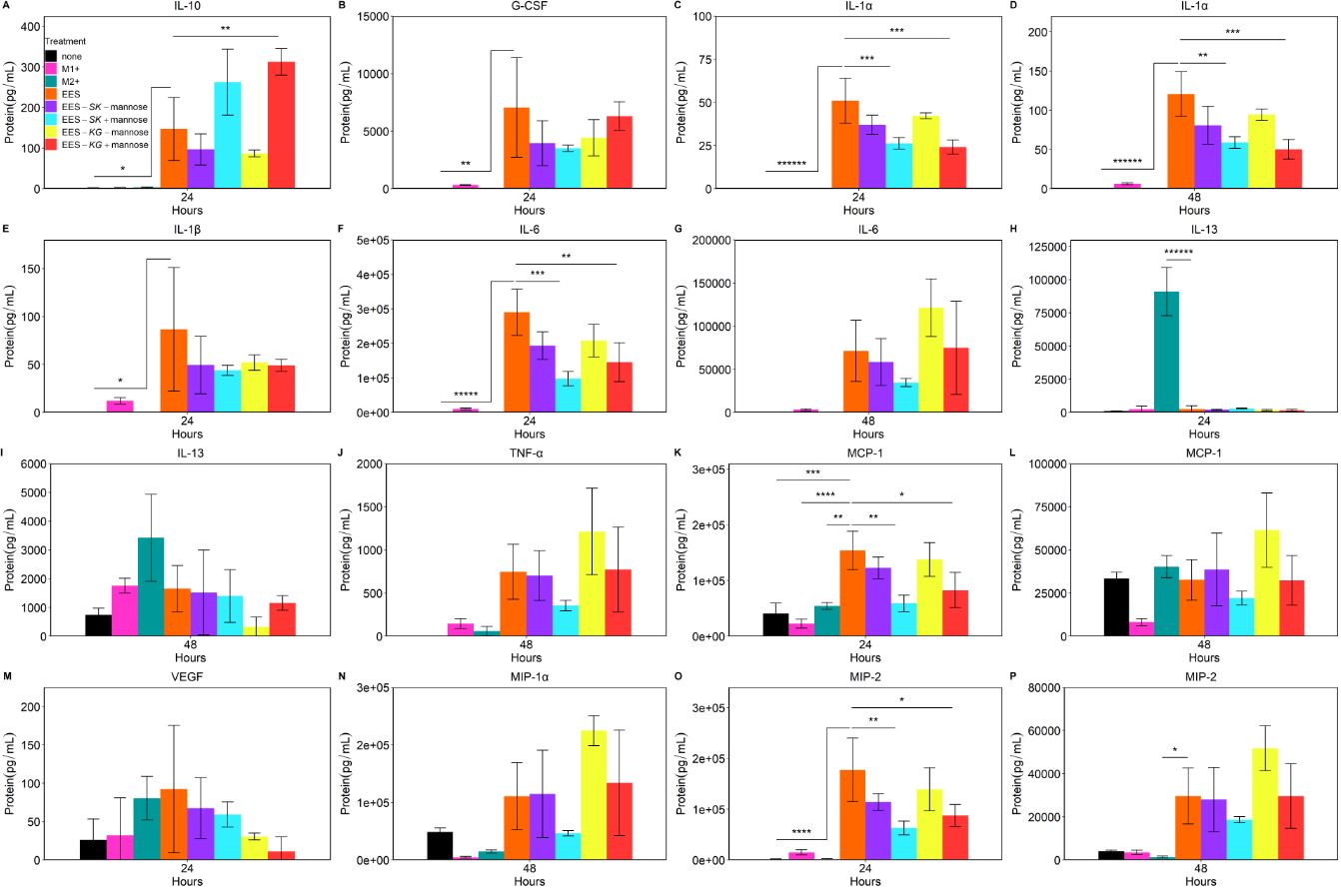

**Figure S6. Remainder of Luminex cytokine profiling assay characterizing EES impact on J774A.1 cell gene expression** **and related to Figure 6**

Cytokine protein concentration was quantified after J774A.1 cells were treated with nothing (none), LPS and IFN-γ (M1+), IL-4 and IL-13 (M2+), EES, EES-*SK* with and without mannose (EES-*SK* -mannose, EES-*SK­* +mannose) and EES-*KG* with and without mannose (EES-*KG* -mannose, EES-*KG* +mannose) at 24 and 48 hr post initial treatment. Data is mean ± SD; *p<0.05, **p<0.01, ***p<0.001, ****p<0.0001, *****p<0.00001, ******p<0.000001. Significance shown is comparing EES condition to all other conditions.

**Supplemental items**

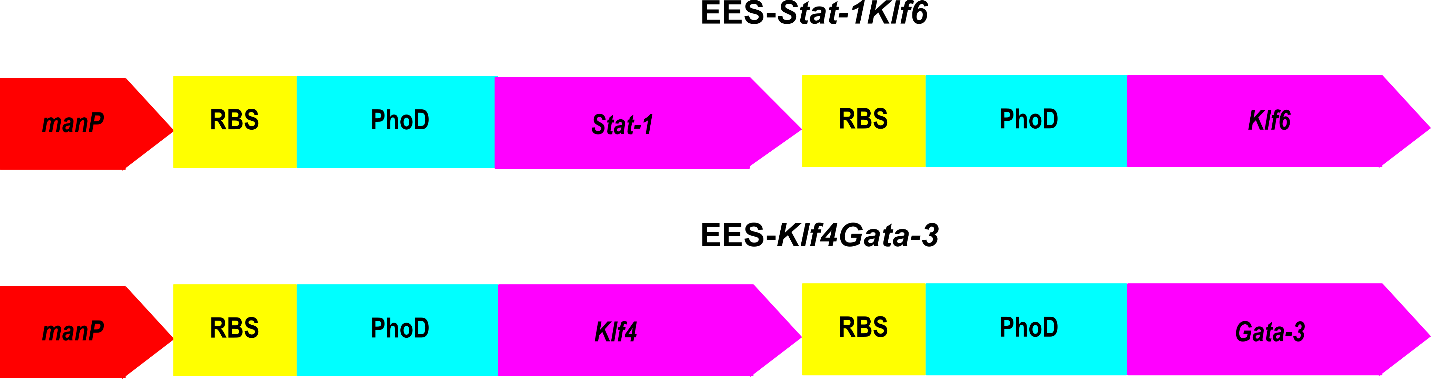

Figure S7: Visualization of polarization operons created for EES expression

The operons for polarization were designed to be transcriptionally controlled by the mannose promoter (*manP*). Then a Gram-positive ribosomal binding site (RBS) and secretion peptide (PhoD) were placed in front of each gene in both operons.
