## Supplementary figures for "Engineered endosymbionts capable of directing mammalian cell gene expression"

| Oligonucleotide name | Sequence | Supplier |
| --- | --- | --- |
| pDR111 Phyper linear F | CTGCATAAAAAACGCCCGGC | IDT |
| pDR111 Phyper linear R | TAAGCTTagtcgacagctagccg | IDT |
| pDR111 lacI linear F | cggctagctgtcgactAAGCTTA | IDT |
| pDR111 lacI linear R | TGAGTTAGGATCCTGAGCGCC | IDT |
| Pman F | GCCGGGCGTTTTTTATGCAGgggattattccttgcttttttt | IDT |
| Pman R | cggctagctgtcgactAAGCTTAaggaaacctcctttaaagtgt | IDT |
| Mannose regulator F | TAAGCTTagtcgacagctagccgattgctgagagagaatggcc | IDT |
| Mannose regulator R | GGCGCTCAGGATCCTAACTCAttaataatccaaatgagataaaaat | IDT |
| Bgal F NLS | CCTAAAAAAAAAAGAAAAGTGACCATGATTACGGATTCACTG | IDT |
| Bgal F no NLS | ttgcgcaatcagtgggagcgACCATGATTACGGATTCACTG | IDT |
| Bgal IPTG R | CCGAATTAGCTTGCATGcggTTATTTTTGACACCAGACCAACT | IDT |
| Bgal mannose R | cattctctctcagcaatcggTTATTTTTGACACCAGACCAACT | IDT |
| SbfI mannose F | gaggaagcggCCTGCAGGaggaaacctcctttaaagtgtgaa | IDT |
| SbfI mannose R | gaggaagcggCCTGCAGGagtcgacagctagccgattg | IDT |
| Stat-1 F | tttcctTAAGCTTagtcgacAAGGAGGTAAGTTatggcgt | IDT |
| Stat-1 R | CAGCTTTTCGCCCACcggccgttacacggtgctcatcatgc |  |
| Klf6 F | caccgtgtaacggccgGTGGGCGAAAAGCTGAAGGA | IDT |
| Klf6 R | AGCTTGCATGcggctagcTTACAGATGCCTTTTCATGTGGAG | IDT |
| Klf4 F | tttcctCCTGCAGGagtcgacagAAGGAGGTAAGTTat | IDT |
| Gata-3 F | ACGACATTTCTAAcggccgGTGGGCGAAAAGCTGAAGGA | IDT |
| Gata-3 R | AGCTTGCATGcggctagcTTAGCCCATAGCAGTAACCA | IDT |
| Bgal NLS Gblock | CAATTAAGCTTagtcgacagAAGGAGGTAAGTTatggcgtatgactcaagatttgacgaatgggtgcaaaaactgaaagaagaatcatttcaaaacaacacatttgacagaagaaaatttattcaaggagcgggaaaaattgcgggactgtcactgggactgacaattgcgcaatcagtgggagcgCCTAAAAAAAAAAGAAAAGTG | IDT |
| Bgal no NLS Gblock | CAATTAAGCTTagtcgacagAAGGAGGTAAGTTatggcgtatgactcaagatttgacgaatgggtgcaaaaactgaaagaagaatcatttcaaaacaacacatttgacagaagaaaatttattcaaggagcgggaaaaattgcgggactgtcactgggactgacaattgcgcaatcagtgggagcg | IDT |
| Bgal mannose NLS | tcctTAAGCTTagtcgacagAAGGAGGTAAGTTatggcgtatgactcaagatttgacgaatgggtgcaaaaactgaaagaagaatcatttcaaaacaacacatttgacagaagaaaatttattcaaggagcgggaaaaattgcgggactgtcactgggactgacaattgcgcaatcagtgggagcgCCTAAAAAAAAAAGAAAAGTG | IDT |
| Bgal mannose no NLS | tcctTAAGCTTagtcgacagAAGGAGGTAAGTTatggcgtatgactcaagatttgacgaatgggtgcaaaaactgaaagaagaatcatttcaaaacaacacatttgacagaagaaaatttattcaaggagcgggaaaaattgcgggactgtcactgggactgacaattgcgcaatcagtgggagcg | IDT |
| Stat-1 custom gene sequence | AAGGAGGTAAGTTatggcgtatgactcaagatttgacgaatgggtgcaaaaactgaaagaagaatcatttcaaaacaacacatttgacagaagaaaatttattcaaggagcgggaaaaattgcgggactgtcactgggactgacaattgcgcaatcagtgggagcgatgagccagtggttcgagctgcagcagctggacagcaagttcctggagcaggtgcaccagctgtacgacgacagcttccctatggagatcaggcagtacctggcccagtggctggagaagcaggactgggagcacgccgcctacgacgtgagcttcgccaccatcaggttccacgacctgctgagccagctggacgaccagtacagcaggttcagcctggagaacaacttcctgctgcagcacaacatcaggaagagcaagaggaacctgcaggacaacttccaggaggaccctgtgcagatgagcatgatcatctacaactgcctgaaggaggagaggaagatcctggagaacgcccagaggttcaaccaggcccaggagggcaacatccagaacaccgtgatgctggacaagcagaaggagctggacagcaaggtgaggaacgtgaaggaccaggtgatgtgcatcgagcaggagatcaagaccctggaggagctgcaggacgagtacgacttcaagtgcaagaccagccagaacagggagggcgaggccaacggcgtggccaagagcgaccagaagcaggagcagctgctgctgcacaagatgttcctgatgctggacaacaagaggaaggagatcatccacaagatcagggagctgctgaacagcatcgagctgacccagaacaccctgatcaacgacgagctggtggagtggaagaggaggcagcagagcgcctgcatcggcggccctcctaacgcctgcctggaccagctgcagacctggttcaccatcgtggccgagaccctgcagcagatcaggcagcagctgaagaagctggaggagctggagcagaagttcacctacgagcctgaccctatcaccaagaacaagcaggtgctgagcgacaggaccttcctgctgttccagcagctgatccagagcagcttcgtggtggagaggcagccttgcatgcctacccaccctcagaggcctctggtgctgaagaccggcgtgcagttcaccgtgaagagcaggctgctggtgaagctgcaggagagcaacctgctgaccaaggtgaagtgccacttcgacaaggacgtgaacgagaagaacaccgtgaagggcttcaggaagttcaacatcctgggcacccacaccaaggtgatgaacatggaggagagcaccaacggcagcctggccgccgagctgaggcacctgcagctgaaggagcagaagaacgccggcaacaggaccaacgagggccctctgatcgtgaccgaggagctgcacagcctgagcttcgagacccagctgtgccagcctggcctggtgatcgacctggagaccaccagcctgcctgtggtggtgatcagcaacgtgagccagctgcctagcggctgggccagcatcctgtggtacaacatgctggtgaccgagcctaggaacctgagcttcttcctgaaccctccttgcgcctggtggagccagctgagcgaggtgctgagctggcagttcagcagcgtgaccaagaggggcctgaacgccgaccagctgagcatgctgggcgagaagctgctgggccctaacgccggccctgacggcctgatcccttggaccaggttctgcaaggagaacatcaacgacaagaacttcagcttctggccttggatcgacaccatcctggagctgatcaagaacgacctgctgtgcctgtggaacgacggctgcatcatgggcttcatcagcaaggagagggagagggccctgctgaaggaccagcagcctggcaccttcctgctgaggttcagcgagagcagcagggagggcgccatcaccttcacctgggtggagaggagccagaacggcggcgagcctgacttccacgccgtggagccttacaccaagaaggagctgagcgccgtgaccttccctgacatcatcaggaactacaaggtgatggccgccgagaacatccctgagaaccctctgaagtacctgtaccctaacatcgacaaggaccacgccttcggcaagtactacagcaggcctaaggaggcccctgagcctatggagctggacgaccctaagaggaccggctacatcaagaccgagctgatcagcgtgagcgaggtgcaccctagcaggctgcagaccaccgacaacctgctgcctatgagccctgaggagttcgacgagatgagcaggatcgtgggccctgagttcgacagcatgatgagcaccgtgtaa | IDT; UniProtKB P42225 |
| Klf6 Gblock | gtgggcgaaaagctgAAGGAGGTAAGTTatggcgtatgactcaagatttgacgaatgggtgcaaaaactgaaagaagaatcatttcaaaacaacacatttgacagaagaaaatttattcaaggagcgggaaaaattgcgggactgtcactgggactgacaattgcgcaatcagtgggagcgATGGATGTACTTCCCATGTGCAGTATTTTTCAAGAGCTGCAAATCGTCCATGAAACCGGGTATTTCTCTGCCTTGCCATCCTTGGAGGAGTACTGGCAGCAAACTTGCCTCGAACTCGAGCGGTACCTCCAGTCCGAACCATGTTATGTGTCCGCATCCGAGATTAAGTTTGATTCCCAAGAGGATCTGTGGACTAAGTTCATTTTGGCCAGAGAAAAGAAAGAGGAAAGCGAACTGAAAATAAGTAGTTCTCCACCTGAAGATAGCTTGATCAGTTCCAGTTTCAATTACAATCTCGAAACTAACTCACTTAATTCCGATGTTTCCAGCGAGTCTTCTGACTCATCTGAGGAATTGTCTCCCACCACCAAGTTTACCTCTGACCCCATTGGAGAGGTGCTGGTGAATAGTGGGAATTTGTCTAGCTCTGTGATTTCAACCCCACCAAGTTCCCCCGAGGTAAACCGGGAATCATCCCAACTTTGGGGATGTGGGCCAGGCGACCTCCCTAGTCCCGGCAAAGTGCGGTCAGGCACATCAGGCAAAAGTGGTGACAAAGGCAACGGTGATGCATCACCCGATGGTAGACGCCGCGTTCACCGCTGTCATTTTAATGGTTGCAGGAAAGTATATACTAAGTCTTCTCACTTGAAGGCTCATCAACGAACACATACAGGTGAGAAGCCTTATCGCTGCAGTTGGGAGGGCTGTGAATGGAGGTTTGCACGGAGCGATGAATTGACCAGGCATTTCAGGAAGCATACAGGGGCTAAACCTTTCAAATGTAGCCATTGCGACAGATGCTTCTCTAGGTCAGATCACCTTGCCCTCCACATGAAAAGGCATCTGTAACCGAATTAGCTTGCATGcgg | IDT; UniProtKB O08584 |
| Klf4 Gblock | CAATTAAGCTTagtcgacagAAGGAGGTAAGTTatggcgtatgactcaagatttgacgaatgggtgcaaaaactgaaagaagaatcatttcaaaacaacacatttgacagaagaaaatttattcaaggagcgggaaaaattgcgggactgtcactgggactgacaattgcgcaatcagtgggagcgATGCGACAACCACCTGGGGAGTCAGATATGGCAGTGTCCGATGCCCTGCTGCCTTCATTTTCCACCTTCGCAAGCGGACCAGCAGGACGGGAGAAGACCCTTCGGCCAGCAGGCGCTCCCACCAACCGCTGGAGGGAAGAGCTTAGCCACATGAAGCGATTGCCACCCTTGCCAGGTCGACCCTACGATCTTGCAGCTACCGTCGCCACCGACCTTGAGTCTGGAGGTGCTGGCGCAGCCTGCTCAAGTAACAACCCCGCTCTCCTTGCAAGGAGAGAGACTGAAGAGTTTAATGATTTGCTTGATCTTGATTTTATTTTGTCTAACTCACTGACACACCAAGAAAGTGTTGCTGCAACAGTGACAACAAGCGCAAGCGCTAGTAGTTCCTCTTCTCCCGCAAGCTCTGGCCCAGCCTCTGCACCTTCCACTTGTTCATTCAGTTACCCTATTCGAGCCGGCGGGGACCCTGGGGTGGCAGCCTCAAATACTGGAGGGGGTCTGCTCTATTCTCGCGAGAGCGCACCTCCCCCAACCGCACCATTTAACTTGGCAGATATTAACGACGTATCTCCCAGTGGCGGATTTGTCGCCGAGCTCCTTAGACCTGAGTTGGATCCCGTATACATTCCTCCCCAGCAGCCTCAACCCCCTGGTGGTGGCTTGATGGGGAAATTCGTGCTCAAAGCCAGTCTGACCACCCCAGGTTCCGAGTATAGCAGCCCCAGCGTAATAAGCGTGTCCAAGGGGTCCCCTGACGGAAGCCACCCCGTTGTAGTTGCTCCTTACTCCGGGGGCCCTCCCAGGATGTGCCCCAAGATAAAACAAGAGGCAGTACCCAGCTGTACCGTGAGTCGGAGTCTTGAAGCTCACCTTTCCGCTGGGCCACAGCTTAGCAATGGTCATCGACCAAACACTCACGATTTCCCCCTGGGACGGCAGCTCCCAACCCGAACAACCCCTACCTTGTCACCAGAAGAGTTGCTCAACAGCCGGGATTGCCATCCCGGTCTCCCACTTCCCCCTGGATTCCACCCTCACCCTGGCCCAAATTATCCCCCATTCCTTCCTGACCAAATGCAATCTCAGGTCCCCTCTTTGCACTATCAAGAACTTATGCCCCCAGGCAGTTGCCTCCCAGAGGAACCCAAGCCCAAAAGGGGTCGACGCTCTTGGCCCCGAAAGAGAACTGCCACTCATACATGTGACTACGCAGGTTGCGGGAAGACCTACACTAAAAGCTCTCATCTTAAAGCACATCTCCGCACACACACAGGCGAGAAGCCCTATCACTGTGACTGGGACGGATGTGGATGGAAATTCGCTCGCAGTGATGAACTGACAAGGCACTACAGAAAGCATACTGGTCACAGGCCATTTCAATGCCAGAAATGTGACCGGGCCTTTAGCAGGTCCGACCACTTGGCACTTCATATGAAACGACATTTCTAA gtgggcgaaaagctg | IDT; UniProtKB Q60793 |
| Gata-3 Gblock | gtgggcgaaaagctgAAGGAGGTAAGTTatggcgtatgactcaagatttgacgaatgggtgcaaaaactgaaagaagaatcatttcaaaacaacacatttgacagaagaaaatttattcaaggagcgggaaaaattgcgggactgtcactgggactgacaattgcgcaatcagtgggagcgATGGAAGTGACTGCCGATCAACCACGCTGGGTGAGCCACCATCACCCAGCAGTCCTGAACGGCCAACATCCCGATACCCATCACCCCGGTCTTGGGCACTCATATATGGAGGCACAGTATCCCCTGACAGAAGAAGTTGATGTACTGTTTAATATAGACGGCCAGGGGAATCATGTGCCTAGTTACTATGGTAATTCTGTGCGGGCAACTGTACAGAGATACCCACCAACCCACCATGGCAGCCAAGTCTGTAGGCCTCCTCTTTTGCACGGGTCTTTGCCTTGGCTCGATGGGGGTAAGGCTCTCAGCTCCCATCACACTGCATCCCCTTGGAATCTTTCACCATTTTCAAAAACAAGTATCCACCATGGGTCCCCCGGACCACTGAGTGTCTATCCTCCTGCATCAAGTTCATCACTTGCTGCTGGGCACAGTAGTCCCCATCTCTTCACATTTCCTCCTACCCCTCCCAAAGACGTCAGCCCAGATCCCTCACTTAGTACTCCTGGTTCCGCCGGTTCTGCCCGCCAAGACGAAAAAGAATGTCTCAAATACCAGGTTCAACTTCCCGACAGCATGAAACTTGAGACATCACACAGTAGAGGTAGTATGACCACACTTGGGGGCGCTTCTAGCAGTGCTCATCACCCAATCACCACATACCCACCATATGTTCCTGAGTACTCAAGTGGATTGTTCCCTCCCTCAAGCCTTCTTGGTGGGTCACCAACAGGTTTCGGATGCAAGTCCCGACCTAAAGCAAGGTCCAGCACTGAAGGTAGGGAATGTGTGAACTGTGGCGCTACCAGTACACCACTCTGGAGACGGGATGGAACTGGACATTATCTCTGCAATGCCTGCGGCCTTTACCATAAGATGAATGGGCAAAACCGCCCTCTTATCAAGCCTAAGCGACGACTTTCCGCAGCACGGCGCGCCGGGACCTCCTGCGCCAACTGCCAAACTACAACCACTACACTGTGGCGCAGAAACGCCAACGGTGACCCAGTTTGTAATGCCTGCGGGCTCTACTATAAGCTTCACAATATTAATAGACCCCTGACCATGAAAAAGGAAGGAATACAGACACGCAATAGGAAGATGTCTTCCAAATCCAAAAAGTGTAAGAAGGTGCATGATGCTTTGGAAGACTTCCCTAAAAGCAGTAGTTTCAATCCCGCCGCCCTTAGCCGCCATATGAGTAGTCTTTCTCATATTTCTCCTTTCTCACACTCTTCTCATATGCTGACTACCCCTACACCCATGCACCCACCTTCAGGTCTCAGCTTTGGCCCACATCACCCAAGTAGTATGGTTACTGCTATGGGCTAACCGAATTAGCTTGCATGcgg | IDT; UniProtKB P23772 |
